# Dual-antigen Doggybone^TM^ DNA vaccine induces potent anti-tumor immunity against immunosuppressive oral cancer

**DOI:** 10.64898/2026.01.21.700768

**Authors:** Grace M.Y. Tan, Chuan Wang, Maria M. Meschis, Josephine Buckingham, Kue Peng Lim, Sok Ching Cheong, Gareth J. Thomas, Sungwon Kim, Lisa Caproni, Helen Horton, Christian H. Ottensmeier, Natalia Savelyeva

**Affiliations:** Head and Neck Centre, Institute of Systems, Molecular and Integrative Biology, University of Liverpool, Liverpool, United Kingdom; Cancer Science, Faculty of Medicine, University of Southampton, Tremona Road, Southampton, United Kingdom; Cancer Immunology and Immunotherapy Unit, Cancer Research Malaysia, Selangor, Malaysia; Department of Oral and Maxillofacial Clinical Sciences, Faculty of Dentistry, University Malaya, Kuala Lumpur, Malaysia; Oral Cancer Research and Coordinating Centre (OCRCC), Faculty of Dentistry, University Malaya, Kuala Lumpur, Malaysia; Touchlight Genetics Ltd, Hampton, United Kingdom; The Clatterbridge Cancer Center, Liverpool, United Kingdom

**Keywords:** DNA cancer vaccine, cancer testis antigens, cold tumor, HPV negative head and neck squamous cell carcinoma

## Abstract

Anti-PD1 blockade benefits only a subset of patients with head and neck squamous cell carcinoma (HNSCC), highlighting the need for approaches that overcome tumor immune resistance. Here, using the doggybone DNA (dbDNA^TM^) platform, we developed CaVac OPT, an optimized dual-antigen DNA vaccine targeting MAGED4B and FJX1, which are overexpressed in most HPV-negative HNSCC and multiple solid tumors. In the MOC-2 oral cancer model, CaVac OPT significantly reduced tumor growth and when combined with anti-PD1 therapy, further delayed progression and improved survival. Immune profiling showed increased infiltration of CD4^+^ and CD8^+^ T cells, expansion of stem-like Tcf1^+^ populations, without increase in regulatory T cells, a reduced M2/M1 macrophage ratio and activation of interferon gamma associated pathways with suppression of tumor-promoting signals. These findings demonstrate that CaVac OPT reprograms the tumor microenvironment, converting cold HNSCC into T cell-inflamed responsive tumors. CaVac OPT represents a promising strategy for achieving durable control of aggressive, immunotherapy-resistant head and neck cancer.

## Introduction

Therapeutic cancer vaccines are increasingly recognized for their potential to enhance clinical outcomes when combined with immune checkpoint inhibitors (ICIs), particularly in patients refractory to standalone immunotherapy^1–3^. These vaccines are engineered to elicit tumor-specific T-cell responses, promoting their infiltration into the tumor microenvironment (TME) and augmenting the efficacy ICIs. Among emerging platforms, DNA vaccines have shown considerable promise. For instance, the DNA vaccine EVX-02, administered alongside ICIs, induced robust T-cell responses in all 10 stage III/IV melanoma patients evaluated, with no disease recurrence observed after one year, a notable contrast to the historical recurrence rates of approximately 30% for nivolumab monotherapy^3^. Similarly, the DNA vaccine SCIB1 targeting the unmutated melanoma antigen gp100, achieved an 85% objective response rate (ORR) in previously untreated, unresectable stage III/IV patients when combined with nivolumab and ipilimumab, with tumor volume reductions ranging from 40 to 100%, by 25 weeks post-treatment^4^. In a phase I/II trial involving advanced hepatocellular carcinoma, combining DNA vaccines targeting neoepitopes with ICI pembrolizumab yielded an ORR exceeding 30%, including an 8% complete response (CR) rate, outperforming the 12-18% response rates typically observed with anti-PD1 monotherapy. Notably, nearly 90% of the patients showed neoantigen-specific T-cell responses, underscoring the capacity of DNA vaccines to generate broad, durable cytotoxic CD8^+^ T cell responses^5^.

DNA vaccines offer distinct advantages, including cost effectiveness, safety, stability and flexibility in design and construction. These attributes enable the construction of complex vaccines incorporating multivalent antigens and immunostimulatory sequences to optimize T-cell induction^6–8^. Similarly, DNA vaccines can encode numerous neoepitopes tailored to a patient’s mutanome, thereby enhancing the induction of tumor-reactive T cells and priming *de novo* responses alongside the expansion of pre-existing T-cell epitopes^5^ Recent studies also suggest that delivery methods, such as *in vivo* electroporation and needle-free injection systems (NFIS), enhance T-cell responses and clinical efficacy^9–11^, offering practical solutions for administration without complex formulations in late-stage cancer patients. However, most current DNA vaccines rely on plasmid DNA (pDNA) produced in bacteria, requiring on antibiotic resistance genes for propagation and necessitating endotoxin removal. While these extraneous bacterial sequences may boost immunogenicity, they also carry risks, such as transgene silencing and the potential transfer of antibiotic resistance to the host’s microbiota^12,13^. This provides a rational to explore enzymatically produced DNA vaccine, such as doggybone DNA (dbDNA^TM^).

dbDNA^TM^ comprises linear, double-stranded DNA with covalently closed ends, synthesized enzymatically via rolling circle amplification by the Phi29 phage polymerase and subsequent cleavage by the N15 phage protelomerase. Unlike pDNA, dbDNA^TM^ is minimalistic, encoding only the immunogen of interest under a eukaryotic promoter with a polyadenylation signal^14,15^. Our previous work demonstrated that dbDNA^TM^ induces cytotoxic T-cell and humoral immune responses comparable to pDNA vaccines^16^ with emerging evidence suggesting it may offer even greater immunogenicity^17,18^. While dbDNA^TM^ has been investigated across various applications, its potential as a therapeutic cancer vaccine remains unexplored, particularly in the context of overcoming an immunosuppressive tumor microenvironment (TME). In this study, we report the optimization of dbDNA^TM^ constructs targeting cancer-testis antigens MAGED4B and FJX1 frequently expressed in solid malignancies, incorporating modifications to enhance T-cell responses. We demonstrate that vaccine candidate CaVac OPT elicits significant, durable protective anti-tumor T-cell response against an HPV-negative oral carcinoma model, characterized by a suppressive TME and which is unresponsive to anti-PD1 ICI. Furthermore, we show that CaVac OPT promotes a shift from M2 to M1 macrophages, underscoring its therapeutic potential.

## Results

### Optimizations to the dbDNA^TM^ vaccine constructs to enhance specific antigen delivery

Here we chose to target cancer-testis antigens MAGED4B and FJX1 due to their prevalent expression across various cancers, including head and neck squamous cell carcinoma (HNSCC) ^8,19,20^, nasopharyngeal carcinoma^21^, lung cancer and other solid tumors^22,23^. Initially, we incorporated several regulatory elements that were previously shown to enhance immunogenicity of plasmid-based DNA vaccines^24^. These included the SV40 enhancer which is designed to augment transcription and promote nuclear translocation, and type C CpG oligodeoxynucleotides (CpG-C) to activate innate immune signaling via TLR9^25,26^ (Supplementary Fig 1). Antigen-specific T cell responses were assessed by interferon gamma (IFN-γ) ELISPOT assays, with comparisons made to the unmodified dbDNA^TM^ constructs, termed Basic 0, which encoded a fusion of each antigen to domain 1 (DOM) of tetanus toxin. The DOM sequence has been shown to enhance induction of CD8^+^ T cells to cancer antigens, resulting in clinical benefits in patients with solid cancers^7,27^. Despite the incorporation of these enhancer elements, none significantly improve T cell responses compared to the Basic 0 construct (Supplementary Fig 1).

To further improve the vaccines performance, we next explored immune-enhancing elements to improve antigen delivery to dendritic cells and refined the antigen design. This included fusion of immune-targeting molecules such as Fms-related tyrosine kinase 3 ligand (Flt3L), or Macrophage Inflammatory Protein-1 alpha (MIP-1a). Both approaches have been tested clinically with demonstration of robust T-cell responses resulting in meaningful clinical responses^11,28,29^. Further variants for FJX1 included engineered conserved consensus antigenic sequences (Con) to break tolerance to this highly conserved antigen^30^. For MAGED4B, we additionally generated variants with a complete and partial deletion of the MAGE homology domain (HD) dHD1 and dHD2 respectively, to eliminate epitopes with potential cross-reactivity to non-target peptides, thereby reducing the risk of off-target cytotoxicity^31,32^. Vaccine variants were compared to a fusion free (FF) construct using a prime/ boost immunization. Among MAGED4B variants, only the dHD1 construct elicited a notably enhanced IFN-γ response (Figure 1A). In contrast, for FJX1, the construct fused to Flt3L induced significantly higher T cell responses (Figure 1B).

**FIGURE 1.**
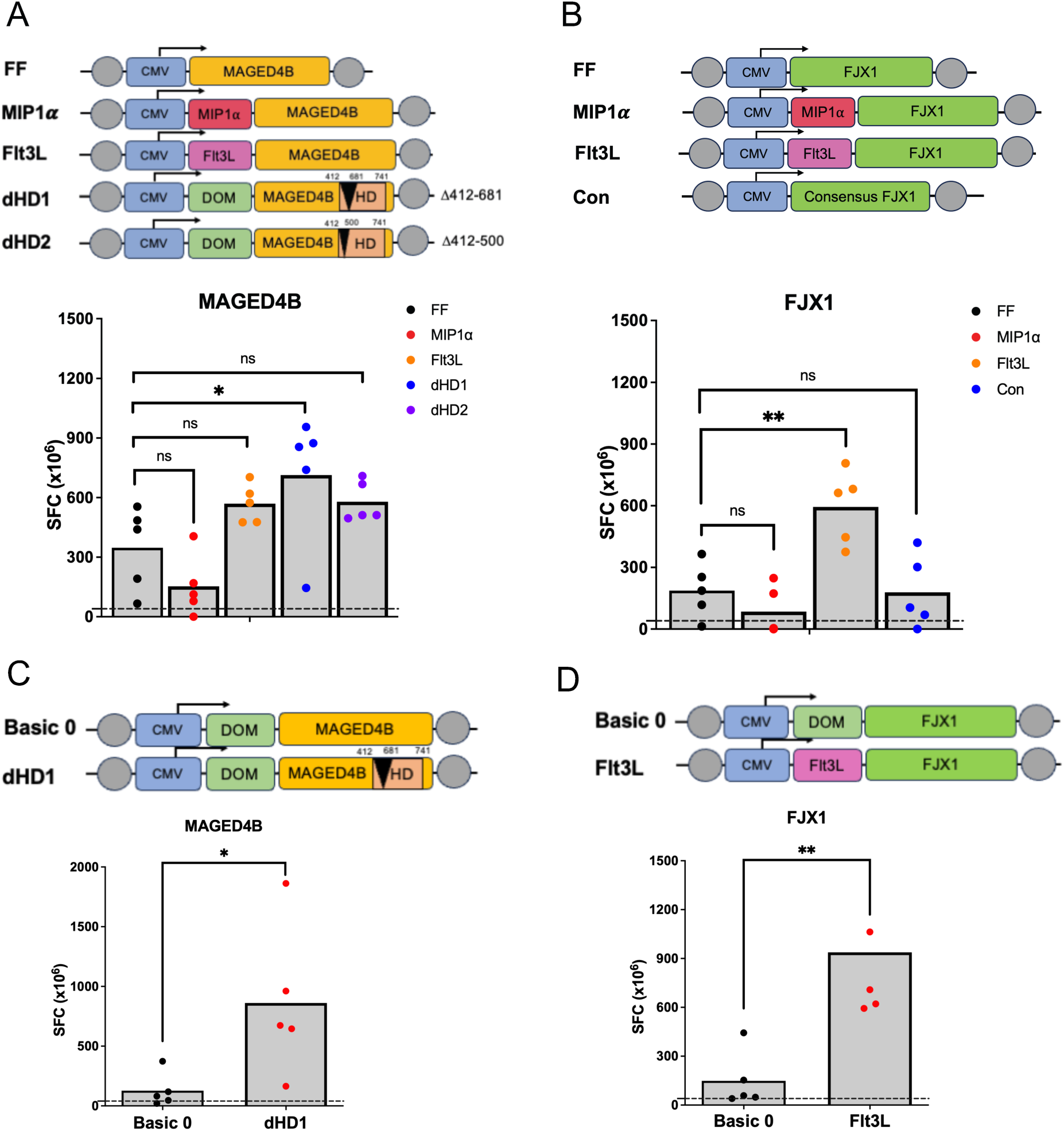
dbDNA^TM^ vaccines encoding MAGED4B and FJX1 elicit antigen-specific T cell responses. C57BL/6 mice (9-12 weeks old) were vaccinated intramuscularly (i.m.) with dbDNA^TM^ vaccine constructs in the anterior tibialis muscle with electroporation. A prime-boost regimen was used, with vaccinations on day one and day twenty-one. Splenocytes were harvested on day thirty post vaccination and stimulated for 40 h with 1μM MAGED4B or FJX1 overlapping peptide pools. Data are presented as spot-forming cells per million (SFC x 10^6^ cells), with background (<10 SFC/10^6^) subtracted. (**A**) Schematic illustration of the dbDNA^TM^ vaccine construct targeting MAGED4B. Each construct as indicated also included the CMV promoter and the SV40 polyA signal. Grey circles represent teRlL sequences, the recognition and cleavage sites for protelomerase TelN. MAGED4B were fused to immune-stimulatory molecules MIP-1α, Flt3L or modified by deletion of the MAGE homology domain (dHD1: amino acids 412-681; dHD2: amino acids 412-500). Vaccine responses were compared to the fusion-free (FF) vaccine. **(B)** FJX1 were fused to immune-stimulatory molecules MIP-1α or Flt3L; the Consensus construct (Con) incorporates consensus sequence modifications derived from FJX1. Vaccine responses were compared to the fusion-free (FF) FJX1 dbDNA vaccines. Statistical comparisons were performed using one-way ANOVA with Tukey’s post-hoc test. Simplified schematic of selected vaccine variants (Basic 0, dHD1 and Flt3L) of **(C)** MAGED4B and (**D**) FJX1 vaccine variants. T cell responses to the most immunogenic variants (dHD1 for MAGED4B; Flt3L for FJX1) were compared to the tetanus DOM fragment-fused vaccine (Basic 0). Statistical analysis determined by unpaired two tailed t-test. Statistical significance is indicated as follows: *p<0.05, **p<0.01, ***p<0.001, ns: not significant.

We next compared the top-performing variants (dHD1 for MAGED4B, and fusion to Flt3L for FJX1) to the Basic 0 constructs. dHD1 induced the strongest IFN-γ responses (Figure 1C) while Flt3L-FJX1 elicited significantly higher responses than Basic 0 (Figure 1D). To assess potential vaccine interference, we compared co-administration of both MAGED4B and FJX1 vaccines at a single site versus opposite flanks and found comparable T cell responses with no evidence of immune interference *in vivo* (Supplementary Fig 2). The combination of dHD1 and Flt3L-FJX1, hereafter referred to as CaVac OPT, was selected for further preclinical evaluation in an oral carcinoma model due to its superior immunogenicity targeting both MAGED4B and FJX1.

### CaVac OPT controls tumor growth and potentiates anti PD-1 therapy

Murine oral carcinoma-2 (MOC-2) is an aggressive head and neck cancer model derived from oral squamous cancerous epithelia^33^. MOC-2 tumors are characterized by a low infiltration of CD8^+^ T cells, low expression of MHC-I, high levels of FoxP3^+^ T regulatory (Tregs) cells and M2 macrophages^33,34^; closely resembling the human oral squamous cell carcinoma (OSCC). Here, MOC-2 cells were engineered to stably express the MAGED4B and FJX1 antigens and were used to evaluate the efficacy of combination dbDNA^TM^ vaccines in controlling tumor growth. Mice were challenged with MOC-2 tumor cells and vaccinated on day three post inoculation with either Basic 0 constructs of MAGED4B and FJX1 or CaVac OPT at low (4µg/mouse) or high (10µg/mouse) doses. Mice that received the vector control exhibited a progressive increase in tumor burden, whereas both vaccines reduced tumor growth at high dose; however, CaVac OPT also induced a significant inhibition of tumor growth at the low dose (p<0.0005) (Figure 2A). Given its superior efficacy at both doses, we further assessed the therapeutic efficacy of CaVac OPT in mice bearing larger MOC-2 tumors. Mice were vaccinated with CaVac OPT on day seven post-tumor inoculation. Compared to controls, vaccinated mice displayed a significantly reduced tumor burden (p<0.05) (Figure 2B).

**FIGURE 2.**
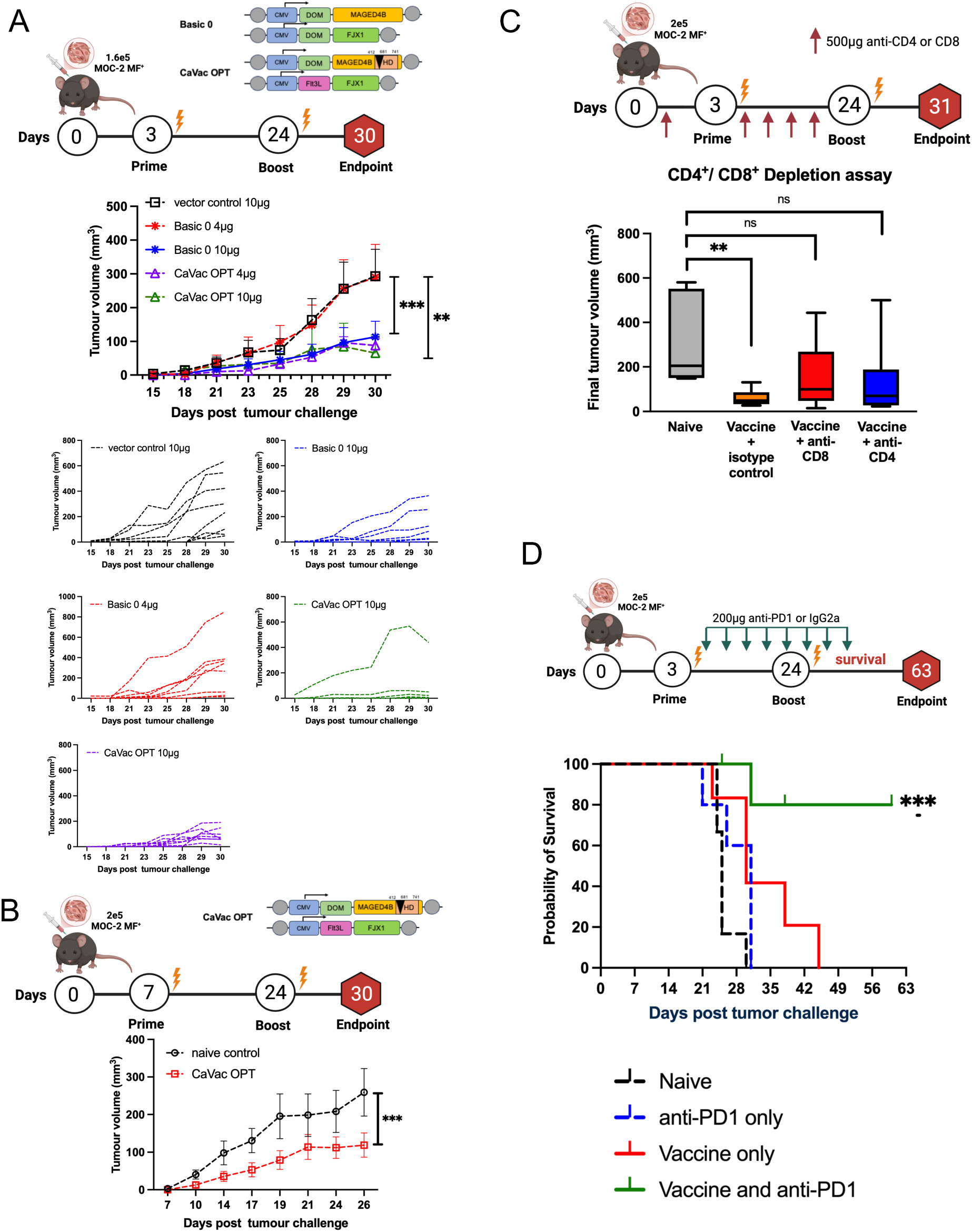
Basic 0 and CaVac OPT vaccine induces superior anti-tumor immunity in the MOC-2 murine oral carcinoma model. (**A**) Schematics of the vaccination schedule: mice were inoculated with 1.6 x 10⁵ MOC-2 tumor cells and vaccinated on day three and day twenty-four post inoculation. Two vaccine combinations were tested: **Basic 0** (Basic 0 MAGED4B, Basic 0 FJX1) and **CaVac OPT** (dHD1 and Flt3L-FJX1). Each combination was administered at low (4μg/mice) and high (10μg/mice) doses and compared to a vector control group. Significant tumor growth reduction was observed in mice received high dose Basic 0 (blue, p<0.01) and low dose CaVac OPT (purple, p<0.001) compared to control. Data (top) are shown as mean ± SEM (n= 8 per group); individual tumor measurements are displayed below. Statistical analysis determined by Dunnett’s multiple comparisons test. (**B**) Experimental setup: mice were challenged with 2 x 10⁵ MOC-2 tumor cells and vaccinated on day seven and day twenty-eight post-inoculation. Tumor volume was measured every three days until day thirty. Data are presented as mean ± SEM (n=8). Statistical significance was determined by the Mann-Whitney test. (**C**) Role of CD4^+^ and CD8^+^ T cells in CaVac OPT-mediated protection. Mice were treated with depleting antibodies against CD4^+^ or CD8^+^ (500 μg /mice, i.p., every 3-4 days in five doses) followed vaccination with CaVac OPT. Final tumor volumes on day thirty-one were compared to mice received antibody isotype control. Statistical analysis determined by Dunnett’s multiple comparisons test. (**D**) Combination of CaVac OPT and anti-PD1 therapy. Mice were challenged with 2 x 10^5^ MOC-2 tumor cells and vaccinated with 4μg CaVac OPT, with anti-PD1 antibody or anti-PD1 antibody alone (200 μg /mice, clone RMP1-14, every 3-4 days from day five). IgG2a isotype control antibody (200 μg /mice, clone 1-1) was used as control. Kaplan-Meier survival curves of each treatment group (Log-rank test). Statistical significance is indicated as follows: *p<0.05, **p<0.01, **3*p<0.001, ns: not significant.

To assess the roles of CD4^+^ and CD8^+^ T cells in CaVac OPT-mediated anti-tumor immunity, we administered anti-CD4 or anti-CD8 depleting antibody to MOC-2 tumor-bearing mice vaccinated with CaVac OPT (Figure 2C) and compared the tumor volumes at the end of experiments. In CaVac OPT-vaccinated mice, depletion of either CD4^+^ or CD8^+^ T cell subsets contributed to loss of protection against tumor challenge, resulting in a higher tumor burden compared to vaccinated mice given an isotype control (Figure 2C). This indicates that both T cell subsets contribute to protection against tumor challenge in this model.

We next evaluated the combination of CaVac OPT with anti-PD1 therapy. Mice were challenged with high tumor burden and vaccinated with CaVac OPT on day three and day twenty-four post-tumor challenge. Starting on day eight post-tumor challenge, mice received eight doses of anti-PD1 or isotype control antibody (Figure 2D). An anti-PD1 alone group was included for comparison, as MOC-2 tumors are known to be resistant to anti-PD1 treatment^35^. The CaVac OPT vaccine significantly sensitized the MOC-2 model to ICI blockade, reducing tumor burden in mice and acting synergistically with anti-PD1. Kaplan-Meier analysis further confirmed that combination therapy significantly extended the mean survival compared to naïve mice, vaccine alone or anti-PD1 alone (Figure 2D). These findings demonstrate that combining CaVac OPT with anti-PD1 enhances anti-tumor immunity in the ‘cold’ MOC-2 tumor model, conferring protection against the disease.

### CaVaC OPT facilitated tumor infiltration by T cells

We next utilized immunohistochemistry (IHC) to characterize T cell infiltration within TME. MOC-2 tumors were harvested at two time points following either priming alone (harvest at day fifteen) (Figure 3A) or after prime and boost vaccinations (harvest at day thirty post-inoculation) (Figure 3B). IHC analysis at day fifteen (post prime) revealed a notable infiltration of CD8^+^ and CD4^+^ T cells in the CaVac OPT-treated group, without an increase in FoxP3^+^ Tregs (Figure 3C). This led to a significant increase in CD8^+^/CD4^+^ T cell ratio compared to the controls (data not shown), with no significant change in the CD8^+^/FoxP3^+^ ratio (Figure 3E). By day thirty, there was a significant absolute increase in antigen-specific CD8^+^ and CD4^+^ T cells, but not FoxP3^+^ T cells compared to unvaccinated controls (Figure 3D), also resulting in a significant increase in the ratio of CD8^+^/FoxP3^+^ Treg ratios (Figure 3F).

**FIGURE 3.**
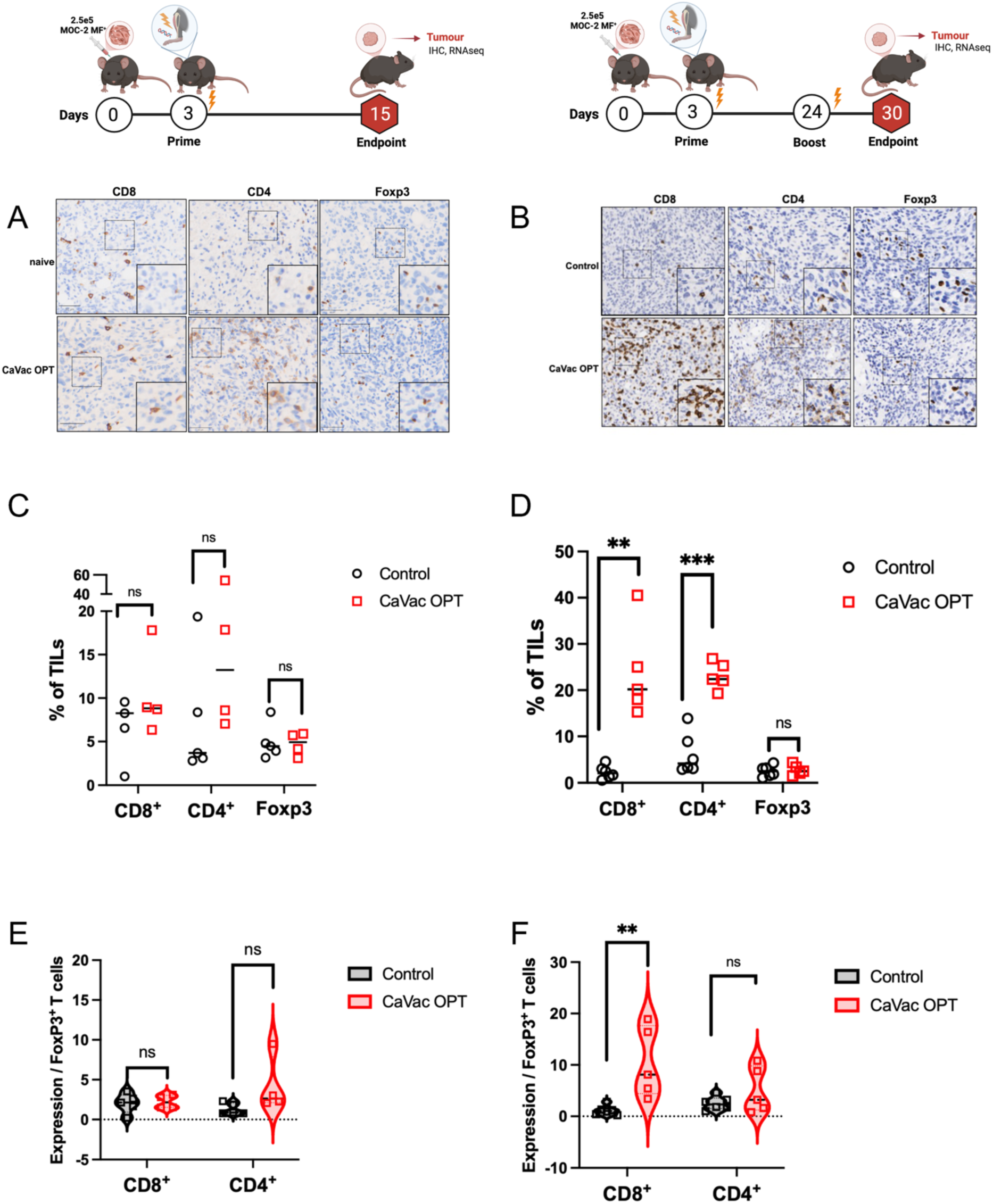
CaVac OPT vaccination enhances anti-tumor immunity via intra-tumoral T cell infiltration of MOC-2 tumors. C57BL/6 mice were subcutaneously injected with 2.5 x 10⁵ MOC-2 tumor cells and either treated or left untreated. Tumors were excised, fixed in 4% paraformaldehyde and processed for immunohistochemistry (IHC) analysis. (**A–B**) Representative IHC images of tumor infiltrating T cells following **(A)** prime vaccination alone on day three post tumor inoculation or **(B)** prime and boost vaccination administered on day three and day twenty-four post tumor inoculation. Tumor sections from control and CaVac OPT-treated mice were stained for CD8⁺, CD4⁺, and FoxP3⁺ cells. Positive staining is indicated in brown. Scale bar = 20 μm. Statistical significance is indicated as follows: *p<0.05, **p<0.01***p<0.001, ns: not significant. % of CD8⁺, CD4⁺, or FoxP3⁺ for immunohistochemistry staining are presented in scatter plots. Ten random fields from each section were counted. (**C)** Dot plots depicting the percentage of CD8⁺, CD4⁺, and FoxP3⁺ cells in across control mice (n=5) and vaccinated mice (n=4) following prime vaccination. (**D**) Dot plots depicting the percentage of CD8⁺, CD4⁺, and FoxP3⁺ cells in control mice (n=5) and vaccinated mice (n=6) following prime and boost vaccination. (**E**) Violin plots illustrate the ratios of CD8⁺/FoxP3⁺ and CD4⁺/FoxP3⁺ T cells across control mice (n=5) and vaccinated mice (n=4) after prime vaccination. **(F)** Violin plots illustrate the ratios of CD8⁺/FoxP3⁺ and CD4⁺/FoxP3⁺ T cells across treatment groups after prime and boost vaccination. Statistical analysis was performed using the Mann–Whitney test.

### CaVaC OPT vaccination reprograms the transcriptome towards enhanced IFN-γ signaling and cytotoxic T cell recruitment

To assess changes in gene expression, we performed bulk RNAseq analysis on CD45^+^ tumor-infiltrating immune cells to compare the immune landscape between vaccinated and control mice post priming (Figure 4A). We analyzed differentially expressed genes (DEGs) and associated signaling pathways between the vaccinated and control groups. At day fifteen post-vaccination, 23 genes were significantly upregulated and 70 were significantly downregulated (log_2_FC>|0.5|, padj<0.1) (Figure 4B, Supplementary table 1). Notably, genes encoding molecules associated with cytolytic T-cell function such as *Gzmk*, *Ifng* and *Nkg7* were markedly upregulated (log_2_FC +3.0, +2.4, +1.6, respectively; padj<0.1) in the vaccinated group (Figure 4B). Additionally, expression of the T cell homing receptor *Cxcr3* (log_2_FC +1.46) and chemokine *Ccl5* (log_2_FC +1.47) was elevated, indicating enhanced priming and recruitment of cytotoxic lymphocytes to the tumor site. Gene Set enrichment analysis (GSEA) using Hallmark gene sets identified 15 significant enriched pathways (p<0.005), including IFN-γ response (normalized enrichment score [NES] +1.46) pathway and interferon alpha response (NES +1.5) (Figure 4C) (Supplementary table 2). Further enrichment analyses with Reactome and Gene Ontology (GO) databases (Supplementary table 3) highlighted pathways related to T cell receptor complex activation. KEGG pathway analysis (Supplementary table 4) also showed positive enrichment in antigen processing and presentation (mmu04612, NES +2.1). Both DEG and GSEA analyses consistently highlighted enhanced IFN-γ signaling in the vaccinated group (Figure 4D), suggesting infiltration of effector T cells with enhanced IFN-γ signaling and antigen processing and presentation with the vaccinated group (Figure 4D).

**FIGURE 4.**
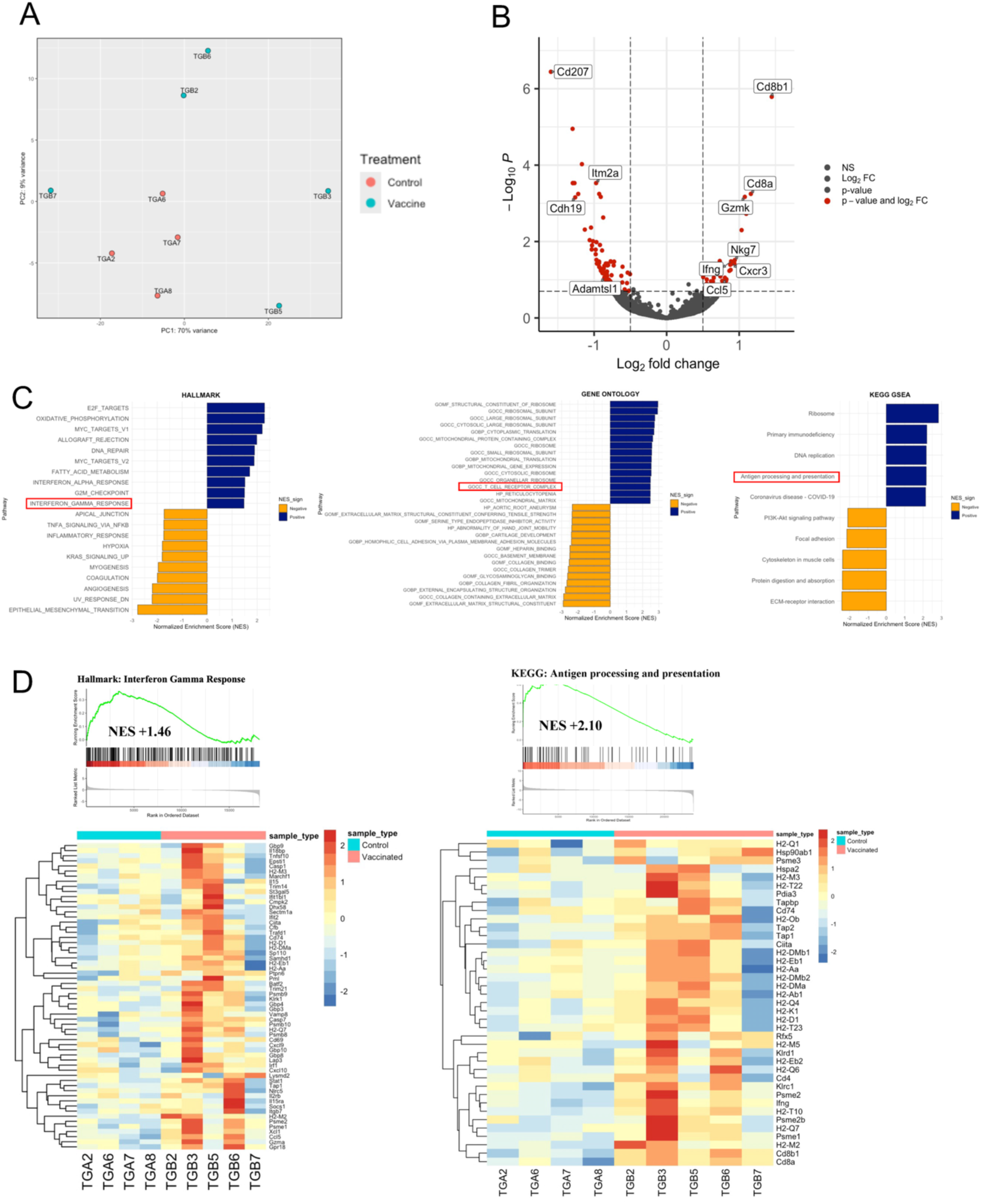
CaVac OPT vaccination reshapes immune cell composition in cold tumors. C57BL/6 mice were subcutaneously injected with 2.5 x 10⁵ MOC-2 tumor cells and treated with or without CaVac OPT. On day fifteen post tumor challenge, tumors were harvested and CD45⁺ tumor-infiltrating lymphocytes (TILs) were isolated for bulk RNA sequencing. **(A)** Principal Component Analysis (PCA) of variance stabilized (VST) bulk RNA sequencing data from CD45^+^ TILs in control (n=4) and CaVac OPT vaccinated samples (n=5). Axes indicate the percentage of variance explained by each principal component. Each dot represents one mouse. **(B)** Volcano plot of differentially expressed genes (DEGs) between control (n = 4) and vaccinated (n = 5) mice. Red dots indicate significantly different genes (log₂FC > 0.5, adjusted p < 0.1). **(C)** Gene set enrichment analysis (GSEA) annotation of the top 15 enriched pathways from Hallmark, top 30 Gene Ontology (GO) and top 10 KEGG databases. **(D)** GSEA ridge plots and heatmaps of interferon-gamma response and antigen processing and presentation gene signatures. Left: Ridge plot and heatmap shows expression profiles of IFN-γ-related genes (NES = 1.46); right: antigen processing and presentation (NES = 2.1) in control and vaccinated tumors.

### CaVac OPT increases CD8^+^ memory T cell differentiation and activation

We next performed bulk RNAseq analysis on CD45^+^ TILs at day thirty post-tumor challenge to assess longitudinal changes in gene expression (Figure 5A). Differential expression analysis identified 512 significant DEGs (padj<0.1, log2FC>0.5) (Supplementary table 5), including 211 upregulated and 301 downregulated genes in vaccinated compared to control mice (Figure 5B). Notably, transcription factor *Tcf7* (log_2_FC +2.55) and *Lef1* (log_2_FC +3.8), both critical transcriptional regulators of memory T cell differentiation, were significantly upregulated. Using the gene set signature-based algorithm ImmucellAI, we observed a significant expansion of CD8^+^ central memory T cells within the T cell population (Figure 5C). Consistent with these findings, IHC analyses of tumor tissues following boost vaccination confirmed a significant enrichment of transcription factor 1 (Tcf1) Tcf1^+^ cells, which exhibited characteristics of T cell memory stem cells (T_SCM_) (Figure 5D). Together, these results suggest that boost vaccination promotes the expansion of CD8^+^ memory T cells, supporting their role in durable anti-tumor protection.

**FIGURE 5.**
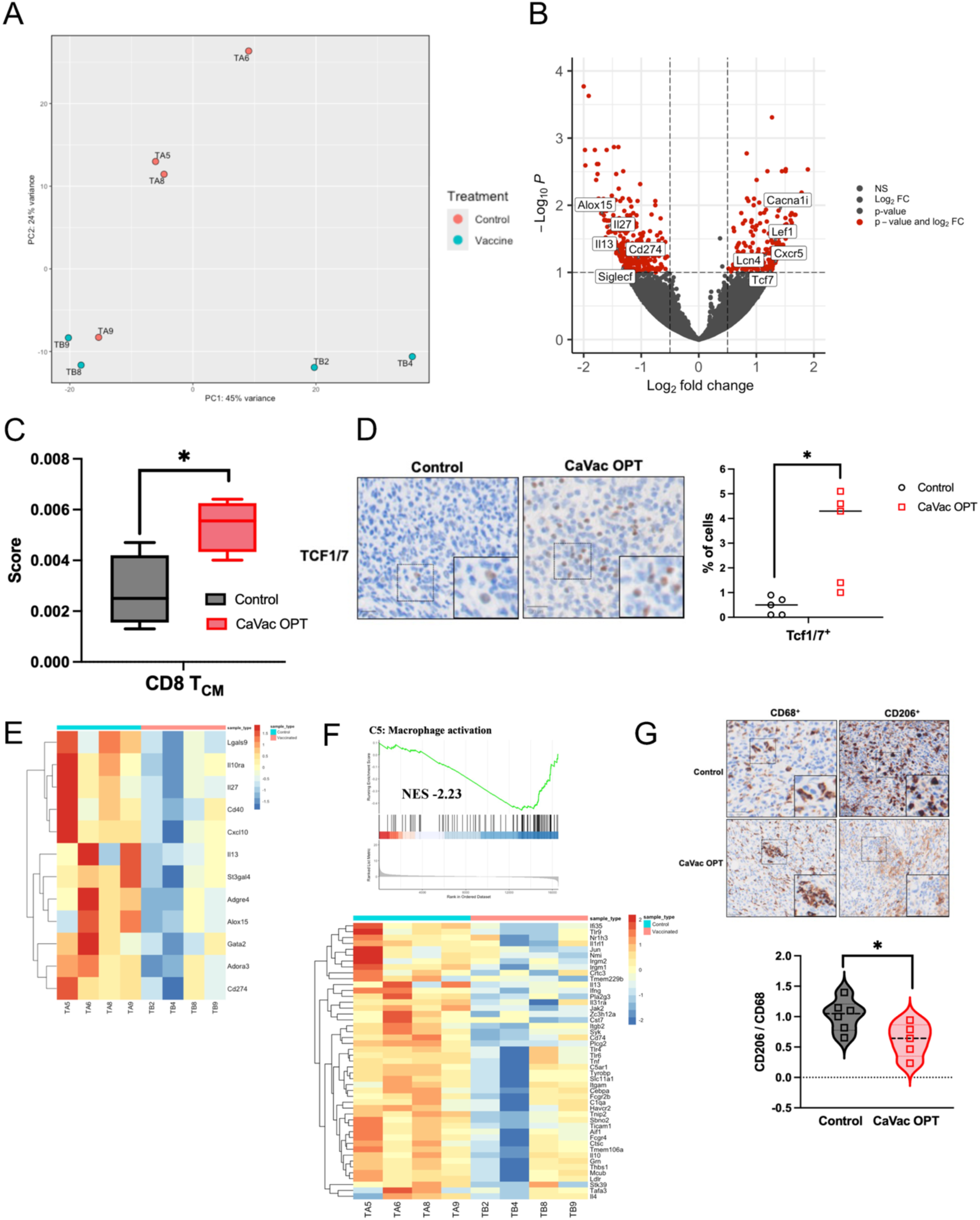
CaVac OPT vaccination promotes memory T cell differentiation and reduces M2-like pro-tumorigenic macrophage in the tumor microenvironment (TME). C57BL/6 mice bearing subcutaneous MOC-2 tumors were treated with or without CaVac OPT. On day thirty post tumor inoculation, tumors were collected and CD45⁺ TILs were isolated for bulk RNA sequencing. Experimental setup: mice were challenged with 2.5 x 10⁵ MOC-2 tumor cells and vaccinated on day three and day twenty-four. Tumor volume was measured every three days until day thirty post tumor challenge. Statistical significance is indicated as follows: *p<0.05, ***p<0.001, ns: not significant. **(A)** PCA of VST bulk RNA sequencing data from CD45^+^ TILs in control and CaVac OPT vaccinated samples. Axes indicate the percentage of variance explained by each principal component. Each dot represents one mouse. (**B**) Volcano plot comparing DEGs between control (n = 4) and vaccinated (n = 4) groups. Red dots denote significant genes (log₂FC > 0.5, adjusted p < 0.1). **(C)** Boxplots illustrating the differences in central memory T cell (T_CM_) estimated by ImmuCellAI between control and vaccinated mice. **(D)** IHC staining of Tcf1^+^ cells in MOC-2⁺ tumors following prime and boost vaccination. Representative images from control and CaVac OPT-treated groups show staining for Tcf1^+^ cells. Brown indicates positive staining. Scale bar = 20 μm. Dot plots show the percentage of Tcf1⁺ cells in vaccinated mice (n = 5) compared with those in the control mice (n = 6). (**E**) Heatmap illustrating expression of macrophage-related DEGs between control and vaccinated tumors. **(F)** GSEA annotation of macrophage activation gene signatures. Ridge plot and heatmap shows gene set enriched in macrophage activation (NES –2.23) in control and vaccinated tumors. **(G)** IHC staining of tumor-infiltrating T cells in MOC-2 tumors following prime and boost vaccination. Representative images from control and CaVac OPT-treated groups show staining for CD68⁺, CD206⁺. Brown indicates positive staining. Scale bar = 20 μm. Violin plots illustrate the ratios of CD206⁺/CD68⁺ across treatment groups. Statistical analysis was performed using the Mann–Whitney test.

### CaVaC OPT vaccine shifts macrophage polarization towards an M1 phenotype in the TME

Macrophages play a critical role in the TME, capable of polarizing toward phenotypes that either promote or suppress tumor progression. Heatmap visualization of bulk RNA-seq data revealed a significant ablation in M2-like gene expression in the vaccinated group (Figure 5E, Supplementary table 5). Specifically, tumor-associated macrophages (TAMs) from vaccinated mice exhibited diminished expression of transcription factor *Gata2* (log_2_FC –1.2, p<0.002) and downregulation of *Il10ra, Il13, Il27* (log_2_FC –1.1, –1.4, –1.6, p<0.001), suggesting that CaVac OPT reduces M2-like macrophage signature in TME. Additionally, expression of inhibitory receptors *Cd274, Adora2a*, and *Adora3a* was attenuated in vaccinated mice (log_2_FC –1.1, –1.0, –1.2, p<0.001), further indicating diminished immunosuppressive activity of M2-like macrophages. The gene *Lgals9*, which encodes galectin-9, was significantly upregulated in control mice compared to vaccinated mice (log_2_FC –0.8, padj<0.1), indicating a higher prevalence of pro-tumorigenic macrophages in the absence of vaccination. GSEA corroborated these findings, identifying negative enrichment of pathways related to regulation of myeloid cell activation in immune response (NES –2.67), macrophage cytokine production (NES –2.3) (Supplementary table 6) and macrophage activation (NES –2.17) in the vaccinated group (Figure 5F).

To validate these transcriptomic signatures at the protein level, we performed IHC analysis on tumor tissue from vaccinated mice. Using CD68^+^ as a pan macrophage marker and CD206^+^ to define the M2-like phenotype^36^, we observed that CaVaC OPT vaccination led to a significant reduction of the CD206^+^/CD68^+^ ratio (p=0.024), as shown in Figure 5G. These findings indicate that CaVaC OPT vaccination effectively shifts macrophage polarization, reducing the proportion of M2-like macrophages within the total macrophage population in mice.

## Discussion

Cancer vaccines are emerging as complementary treatments in tumors with limited responses to ICIs^37^. Recent progress in genetic vaccine platforms, particularly mRNA-based technologies, has reinvigorated interest in DNA-based vaccine approaches. We evaluated a next-generation dbDNA^TM^ vaccine targeting the cancer-testis antigens MAGED4B and FJX1. Although assessed here in a preclinical model of HNSCC, this vaccine is broadly applicable to other solid tumors, as these antigens are widely expressed in other solid tumors^8,19,20^. These antigens represent highly promising targets, enabling the development of effective “off-the-shelf” vaccines without the need for complex epitope discovery or the logistical challenges associated with personalized vaccine production. Moreover, CaVac OPT could serve as a bridging immunotherapy, initiating an early immune response while a personalized vaccine is being developed, thereby potentially enhancing overall clinical benefit. Unlike conventional plasmids, the compact size of dbDNA^TM^ (2 kb smaller than a standard pDNA) may enable efficient nuclear transport, explaining why the SV40 enhancer previously shown to enhance larger DNA constructs^38,39^ did not improve immunogenicity here. Electroporation is likely to bypass the endosomal TLR9 signaling, rendering CpG motifs ineffective, while cytoplasmic pattern recognition receptors, such as cGAS-STING or RIG-I-like receptors, may instead mediate innate sensing of dbDNA^TM16^.

The vaccine constructs were individually optimized for each antigen, resulting in distinct design sequences informed by prior clinical efficacy data. To further refine the dbDNA^TM^ vaccine, we focused on enhancing antigen presentation through strategic modification to the antigenic sequence and incorporation of fusion partners that have proven to be critical for improvement of immunogenicity. Among several fusion partners tested, the tetanus-derived DOM fragment and Flt3L elicited the highest T-cell responses. Unexpectedly, certain fusion partners led to poor T-cell induction in contrast with previous published reports^30,40^. Additionally, removal of the MAGE HD further exposed immunogenic MAGED4B epitopes, suggesting that peripheral tolerance to conserved family domains may limit T-cell induction against MAGED4B.

CaVac OPT vaccination converted cold HNSCC tumors into pro-inflammatory microenvironment, inducing robust CD8^+^ and CD4^+^ T cell responses without an increase in Tregs. Chemokine induction (e.g. Ccl5) and homing receptor expression (Cxcr3) likely facilitate efficient T cell trafficking^41^ into the TME and exert cytotoxic effects through IFN-γ and granzyme K, contributing to tumor cell killing and tumor volume reduction. Following vaccination, infiltrating CD8^+^ T cells exhibited stem-like memory (Tcf7^+^ and Lef1^+^), a subset associated with durable responses^42^ and clinical benefit from PD1 blockade^43,44^, potentially explaining the synergy with anti-PD1 therapy. In parallel, the vaccine reduced the proportion of immunosuppressive M2-like TAMs, which are abundant in HNSCC, where they contribute to tumor progression through immunosuppressive and pro-tumorigenic effects^45–47^. In HNSCC, TAMs are actively recruited to TME and establish direct contact with squamous carcinoma cells. Notably, C-C motif chemokine ligand 18 (CCL18) produced by M2 macrophages has been shown to promote epithelial-mesenchymal transition (EMT) and enhance cancer stemness, thereby driving metastasis^48^. Thus, the ability of our vaccine to reducing M2 polarization may play a crucial role in negatively enriching EMT pathways and enhancing anti-tumor immunity. Together, these findings position CaVac OPT as a rational approach to overcome immune exclusion in HNSCC by simultaneously enhancing T cell priming, memory formation and myeloid repolarization.

In conclusion, CaVac OPT has demonstrated that the dbDNA^TM^ platform represents an immunogenic cancer vaccine capable of eliciting durable, antigen-specific T cell responses. It effectively transforms the immunosuppressive TME through the reduction of M2-like CD206^+^/ CD68^+^ macrophages and results in significant tumor regression. Moreover, it exhibits a synergistic effect when combined with anti-PD1 therapy, underscoring its promise as a platform for future clinical testing.

## Methods

### Generation of dbDNA^TM^ vaccines

All dbDNA^TM^ vaccines were generated and subcloned into the proTLx-K ST^TM^ expression plasmid following previously described protocol^49^. Cancer-tests antigens including Melanoma-associated antigen D4 isoform 1 (MAGED4B, NP_001258991) and human four-jointed box protein 1 (FJX1, NP_055159) were retrieved from the NCBI database. Consensus sequences (Con) were generated by aligning these sequences using the BLOSUM62 algorithm (default matrix) in EMBOSS (European Bioinformatics Institute; EMBOSS Cons Multiple Sequence Alignment EMBL-EBI). Additional sequences, including human macrophage inflammatory protein-1 alpha (huMIP-1a), and FMS-like tyrosine kinase 3 ligand (Flt3L), were obtained from NCBI and optimized for human codon usage with GenSmart^TM^ Codon Optimization software (GenScript). To enhance secretion, the Mus Musculus IgH signal peptide sequence MGWSCIIFFLVATATGVHS was incorporated at the N-terminus of each construct. The DOM fragment (TT_865–1120_)^8^ and the target antigens (MAGED4B or FJX1) were linked using a seven-amino-acid spacer (AAAGPGP), as previously described^8^. Variants of MAGED4B with full (dHD1) or partial homology domain deletions (dHD2) were generated via site-directed mutagenesis. Briefly, plasmid templates contained an expression cassette flanked by dual TelN protelomerase recognition binding sites from *E. coli* phage N15. Templates were denatured with NaOH and quenched in a reaction buffer containing custom primer, dNTPs, Phi29 polymerase and pyrophosphatase. The reaction mixture was incubated at 30°C for 30 or 72 h. Concatemeric DNA was cleaved by adding TelN protelomerase. followed by digestion with 200 units/mL of a template-specific restriction enzyme (New England Biolabs, Hitchin, UK) and 200 units/mL exonuclease *Exo*III (Enzymatics). The digest mixture was purified and precipitated using polyethylene glycol (PEG) 8000.

### *In vivo* mouse vaccination

All mouse-related procedures were approved by the Home Office under project license PP6990832. C57BL/6J mice (6-10 weeks old, 18 to 22 g) were obtained from Charles River laboratories (Kent, UK) and housed in ventilated cages at the University of Liverpool animal facility.

### Tumor cell lines

Mouse oral cancer 2 (MOC-2) cells were cultured in IMDM/F12 (2:1) supplemented with 10 % foetal bovine serum (FBS; Gibco), 100 U/mL penicillin streptomycin (Gibco), 5mg/mL insulin (Sigma, I6634), 400 ng/mL hydrocortisone (Sigma, H0135), 5ng/mL epidermal growth factor (EMD Milipore, 01-107) at 37°C, 5 % CO_2_ in a humidified incubator as previously described. To establish a homogenous MAGED4B– and FJX1-expressing cell line, MOC-2 cells were transduced with retrovirus particles generated using the pBABE-MAGED4B-FJX1-puro vector and HEK293T gag-pol packaging cells (Takara Bio, 631458). The human MAGED4B and FJX1 sequences were cloned into pBABE-Puro vector using NotI and XhoI restriction sites, connected via a GSGSG linker for a single in-frame transcript. Transduced cells were selected with 4 µg/mL puromycin (Gibco) and validated by IHC using anti-MAGED4B (Santa Cruz, sc-393059) and anti-FJX1 (Atlas antibodies, HPA059220) antibodies respectively. Cells were confirmed to be free of mycoplasma and rodent pathogens using PCR analysis (Mouse/Rat CLEAR panel, Charles River) and used at low passage for *in vivo* studies.

### In vivo immunogenicity and efficacy of dbDNA^TM^ vaccines

To evaluate antigen-specific immune responses, mice were randomly assigned to experimental groups and vaccinated intramuscularly (i.m) in both anterior tibialis muscles using electroporation with a custom TriGrid® pulse generator (Ichor medical systems, San Diego, CA). Booster doses were administered 21 days after priming, optimized for eliciting T-cell responses with DNA vaccines.

In tumor-bearing models, mice were subcutaneously (s.c) injected on day 0 with 1.6-2.0 ×10^5^ cells MOC-2 expressing MAGED4B-FJX1^+^. Between day three and seven-post inoculation, mice were randomized to receive one of the following: a control dbGFP vector, combined dbDNA^TM^ of Basic 0 MAGED4B and Basic 0 FJX1, or an optimized formulation (CaVac OPT) dbDNA^TM^ dHD and Flt3L-FJX1. Dosing varied by experimental conditions, and all groups received boosters twenty-one days post-priming. Tumors were measured every three days, and mice were euthanized when tumor volumes exceeded 500mm^3^ or if ulceration reached humane endpoint thresholds. Tumor volume was calculated as 0.5 × (width^2^ x length). Tumor specimens and splenocytes were harvested as required for downstream analysis.

### Ex vivo IFN-γ ELISPOT

Splenocytes were isolated via mechanical dissociation and Lymphocyte density gradient (Stemcell Technologies). Cells were cultured in RPMI supplemented with FBS, P/S, 2mM L-glutamine (Gibco), 1mM sodium pyruvate (Gibco), 2-Mercaptoethanol (Merck) and 1x non-essential amino acids (Gibco). IFN-γ ELISpot assays were performed per BD Biosciences protocols using 2 × 10⁵ splenocytes/well, stimulated with no peptide (background control), 1 µM tetanus p30 (Cambridge Research Biochemistry), or overlapping peptide pools (OPP) of MAGED4B or FJX1 (JPT Peptide, Germany). IFN-γ secreting cells were detected using a biotin-conjugated anti-mouse IFN-γ antibody (BD Biosciences, UK), followed by streptavidin-ALP (Mabtech, UK) and BCIP/NBT substrate (Mabtech, UK). Spot-forming cells (SFCs) were quantified with an automated ELISPOT reader (Aelvis GmbH) and expressed as mean SFCs per 10⁶ cells^16^.

### Combination therapy and T cell subset depletion

To evaluate synergy between vaccination and immune checkpoint inhibition, tumor-bearing mice were assigned to four groups: (1) control vaccine dbGFP, (2) 4µg CaVac OPT+ IgG2a isotype control, (3) 4µg of CaVac OPT + anti-PD1 and (4) anti-PD1 alone. Beginning five days post vaccination, 200µg of anti-PD1 (Clone RMP1-14) or IgG2a isotype control (clone 1-1) was administered intraperitoneally (i.p.) every three or four days in four weeks, for a total of eight doses. For T cell depletion studies, 500 µg of anti-CD4 (clone GK1.5) or anti-CD8 (clone YTS 169) mAbs via i.p. injection, while non-depleted controls received 500 µg of anti-IgG2b isotype control (clone I-1034), for a total of five doses. Depletion was validated via flow cytometry after the fourth administration.

### Tumor Dissociation and tumor infiltrating lymphocytes (TILs) Isolation

Tumors were harvested and processed using the Mouse Tumor Dissociation Kit (Miltenyi Biotec, 130-096-730) and the gentleMACS Dissociator (Miltenyi Biotec) following the manufacturer’s protocol. CD45^+^ TILs were isolated using Mouse CD45 TIL MicroBeads (Miltenyi Biotec, 130-110-618). Biological replicates consisted of tumors harvested from individual mice per treatment group.

### RNA Extraction and Sequencing

RNA samples were extracted from TILs using the RNeasy mini-Kit (QIAGEN). RNA purity, concentration, and integrity were assessed using a NanoDrop spectrophotometer, Qubit 2.0 Fluorometer, and Agilent 2100 Bioanalyzer respectively. Only samples with RNA integrity number (RIN ≥ 4) were used for sequencing. Messenger RNA (mRNA) was enriched from total RNA using poly-T oligo attached magnetic beads (polyA selection). Libraries were prepared, pooled, and sequenced on the Illumina NovaSeq 6000 platform (PE150 mode) (Novogene) to a minimum depth of 6 Gb reads per sample. Raw reads were quality trimmed to remove adapter sequences, reads containing >10% ambiguous bases (N), or low-quality bases. Clean reads were aligned to the *Mus musculus* reference genome (mm39, NCBI RefSeq GCF_000001635.27) using HISAT2 (version 2.0.5)^50^.

### Gene expression and pathway analysis

Gene counts were normalized and analyzed for differential expression analysis using the DESeq2 package (v 1.48.0) in R (v 4.5.0) statistical software^51^. Differentially expressed genes (DEGs) were defined by a fold change (FC) ≥ 1.5 and an adjusted P-value (padj) < 0.1. Principal component analysis (PCA) plot was visualized using the “ggplot2” R package (v 3.5.2)^52^. The clustering analysis of DEGs was carried out using the “tidyheatmaps” package (v 0.2.1) of R^53^. The enrichment of pathways was identified by GSEA method using clusterProfiler (v 4.16.0)^54^. Functional terms were retrieved from the Mouse Signature Database (MSigDB) gene sets R package (v 10.0.2), including the biological process (BP), molecular function (MF), and cellular component (CC) ^55,56^. Then, the analysis was performed to identify significantly different regulatory pathways using the Kyoto Encyclopedia of Genes and Genomes (KEGG), a major public pathway-related database. The results of GSEA were visualised with ggplot2 and enrichplot (v 1.28.0)^57^.

### Immune cell infiltration profiling

Immune cell composition within TILs was inferred using ImmuCellAI-mouse (http://bioinfo.life.hust.edu.cn/ImmuCellAI-mouse/)^58^. Normalized gene expression matrices from treatment were uploaded to predict the relative abundance of immune subsets.

### Immunohistochemistry (IHC)

Tumor samples were fixed in PBS with 4% paraformaldehyde before processing into formalin-fixed paraffin-embedded (FFPE) blocks. Sections of 3 µm were cut and processed on the Leica Bond-RX™ automated staining platform for IHC. Following antigen retrieval and blocking with 10% normal goat serum in PBS (Cell Signaling Technology), tissue sections were incubated with primary antibodies to CD8α (1:100, D4W2Z, Cell Signaling), CD4^+^ (1:100, D7D2X, Cell Signaling), FoxP3^+^ (1:200, D6O8R, Cell Signaling), CD68^+^ (1:600, E3O7V, Cell Signaling), CD206^+^ (1:200, E6T5J, Cell Signaling), TCF1/TCF7 (1:50, C63D9, Cell Signaling). Antigen retrieval was performed using the Bond™ Polymer Refine Detection Kit with either ER1 buffer (10mM sodium citrate buffer) for 20 minutes or ER2 buffer (Tris-EDTA buffer pH 9.0) for 20 minutes (used for FoxP3 staining). Detection was carried out using an HRP-conjugated secondary antibody, followed by DAB chromogenic staining. Sections were counterstained with hematoxylin and mounted for microscopic analysis.

### Quantitative image analysis

Slides were scanned at 20× magnification using PhenoImager Fusion (v2.3.1, Akoya Biosciences). Images were visualized and annotated using QuPath v0.5.111. For each tumor sample, ten regions of interest (ROIs) measuring 500 µm² were selected and analyzed using automated cell detection. Colour deconvolution was applied to separate individual stains, and immune cell populations (CD8^+^, CD4^+^, FoxP3^+^, CD68^+^, CD206^+^, Tcf1/7^+^) were quantified using QuPath’s positive cell detection algorithm. Quantification results were exported and further processed in Microsoft Excel.

### Statistical analysis

Statistical significance was determined using a two-tailed Mann-Whitney test (two-group comparisons), one-way ANOVA (multi-group comparisons), or log-rank Mantel-Cox test (survival analysis) in Prism 10.4.2 (GraphPad, CA, USA). P-values ≤ 0.05 were considered significant (*p < 0.05; **p < 0.01; ***p < 0.001; ****p < 0.0001).

## Acknowledgements

The authors gratefully acknowledge the skilled technical support provided by the Biomedical Services Unit at the University of Liverpool. We are also grateful to the team at Research Histology at the University of Southampton for their expert assistance with immunohistochemical staining.

## Author contributions

C.W. and S.K. conducted experiments related to vaccine design and construction. G.M.Y.T., C.W., and M.M. performed in vivo experiments and data acquisition. G.M.Y.T. analyzed the bulkRNA-seq data. C.W., L.C. and H.H. contributed to design of vaccine candidates. G.J.T contributed to developing methodology. K.P.L., S.C.C., contributed to conceptualization.

C.H.O. contributed to conceptualization and data interpretation. N.S. conceived the ideas, designed the study and supervised the experiments. G.M.Y.T., M.M. and N.S. wrote the manuscript. Every author contributed to the significant revision of the manuscript for essential intellectual content and gave their final approval for the version to be published.

## Data and materials availability

Sequencing data of bulk RNA-seq were deposited at GEO submission with the accession numbers: GSE310571

