## Supplemental figures and tables for "Dual-antigen Doggybone^TM^ DNA vaccine induces potent anti-tumor immunity against immunosuppressive oral cancer"

#### SUPPLEMENTARY FIG 1

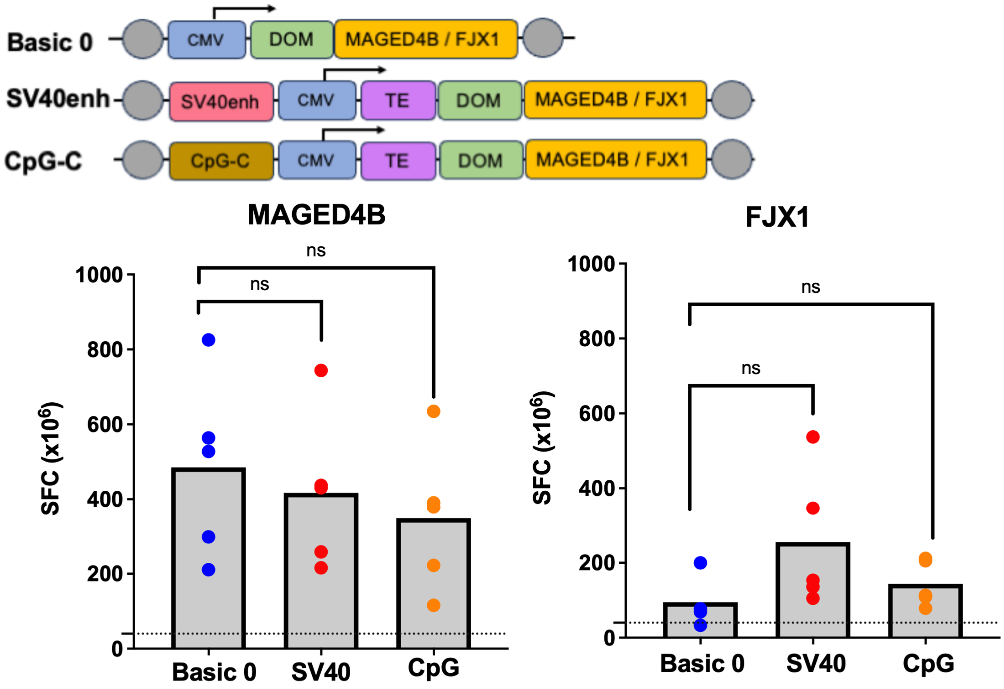


Schematic illustration of the dbDNA$^{TM}$ vaccine construct. C57BL/6 mice (9-12 weeks old) were vaccinated i.m. with dbDNA$^{TM}$ vaccine constructs in the anterior tibialis muscle with electroporation. A prime-boost regimen was used, with vaccinations on day zero and day twenty-one. Splenocytes were harvested on day30 and stimulated for 40 h with 1µM MAGED4B or FJX1 overlapping peptide pools. Data are presented as spot-forming cells per million (SFC x $10^6$ cells), with background (<10 SFC/$10^6$) subtracted. Statistical significance is indicated as ns: not significant. Each construct as indicated also including the CMV promoter and the SV40 polyA signal. Grey circles represent teRlL sequences, the recognition and cleavage sites for protelomerase TelN. regulatory elements SV40enhancer (SV40enh) or type C CpG oligodeoxynucleotides (CpG-C) were included into dbDNA$^{TM}$ vaccines. Vaccine responses were compared to the tetanus toxin fragment C (DOM)-based comparators (Basic 0).

### SUPPLEMENTARY FIG 2

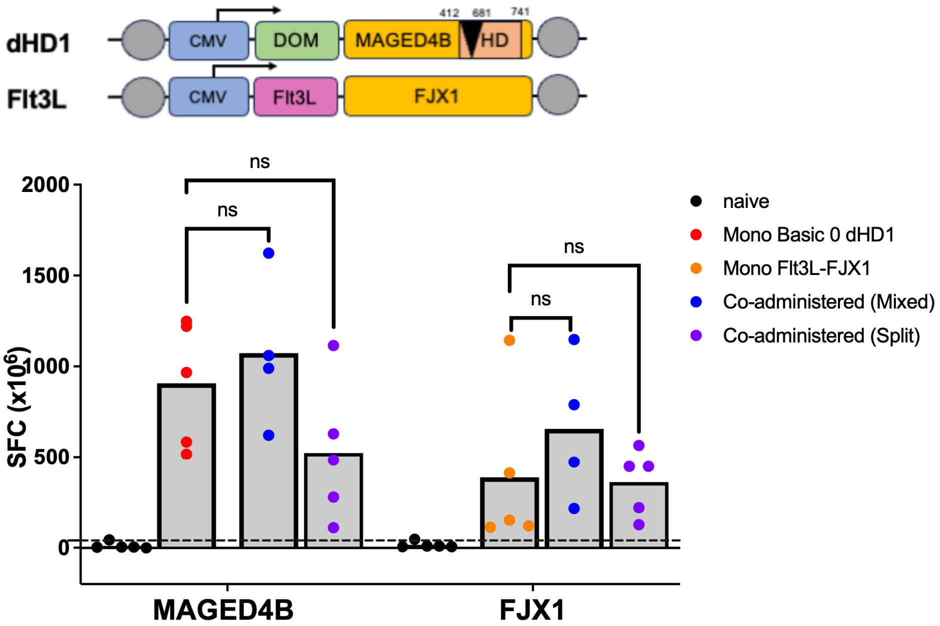


**SUPPLEMENTARY FIG 2:** Schematic illustration of the dbDNA$^{TM}$ vaccine construct tested. C57BL/6 mice (9-12 weeks old) were vaccinated i.m. with dbDNA$^{TM}$ vaccine constructs in the anterior tibialis muscle with electroporation. A prime-boost regimen was used, with vaccinations on d 1 and d 21. Splenocytes were harvested on d 30 and stimulated for 40 h with 1μM MAGED4B or FJX1 overlapping peptide pools. Data are presented as spot-forming cells per million (SFC x $10^6$ cells), with background (<10 SFC/$10^6$) subtracted. Statistical significance is indicated as ns: not significant. Schematic shows combinations of tested vaccine variants: dHD1(MAGED4B) and Flt3L-FJX1. Immune responses from co-administered vaccines mixed at a single site were compared to those administered at separate flanks. Mono-vaccinated controls (Mono basic 0 dHD1, red; Mono Flt3L-FJX1, orange) were included for comparisons.

Supplementary Table 1: Differentially expressed genes (DEGs) at day fifteen post tumor challenge

| | Gene name | log2FC | padj | regulation |
|---|---|---|---|---|
| 1. | Fcgbp | 4.03 | 0.0694 | Up |
| 2. | Trbv29 | 4.02 | 0.0322 | Up |
| 3. | ENSMUSG00000121251 | 3.37 | 0.0883 | Up |
| 4. | Gzmk | 3.05 | 0.0007 | Up |
| 5. | Cd8b1 | 2.59 | 0.0000 | Up |
| 6. | Gm44175 | 2.59 | 0.0376 | Up |
| 7. | Cd8a | 2.54 | 0.0006 | Up |
| 8. | Ifng | 2.41 | 0.0329 | Up |
| 9. | H2-M2 | 2.40 | 0.0815 | Up |
| 10. | Wdr95 | 2.36 | 0.0880 | Up |
| 11. | Gapdh | 2.31 | 0.0050 | Up |
| 12. | Rpl21 | 1.77 | 0.0019 | Up |
| 13. | Ms4a4b | 1.66 | 0.0550 | Up |
| 14. | Nkg7 | 1.55 | 0.0305 | Up |
| 15. | Ccl5 | 1.47 | 0.0968 | Up |
| 16. | Cxcr3 | 1.47 | 0.0376 | Up |
| 17. | Gm8369 | 1.41 | 0.0583 | Up |
| 18. | Baiap3 | 1.31 | 0.0404 | Up |
| 19. | Trim7 | 0.94 | 0.0723 | Up |
| 20. | Klhl36 | 0.81 | 0.0573 | Up |
| 21. | Acsf3 | 0.63 | 0.0976 | Up |
| 22. | Gm6563 | 0.58 | 0.0447 | Up |
| 23. | Ckb | 0.56 | 0.0841 | Up |
| 24. | Homer1 | -0.56 | 0.0710 | Down |
| 25. | Nr1d2 | -0.69 | 0.0454 | Down |
| 26. | Zfp667 | -0.74 | 0.0783 | Down |
| 27. | Mustn1 | -0.87 | 0.0334 | Down |
| 28. | Edil3 | -0.90 | 0.0417 | Down |
| 29. | Rhoj | -0.96 | 0.0329 | Down |
| 30. | Cthrc1 | -1.02 | 0.0756 | Down |
| 31. | Slc5a3 | -1.06 | 0.0023 | Down |
| 32. | Gja1 | -1.07 | 0.0412 | Down |
| 33. | Colgalt2 | -1.08 | 0.0375 | Down |
| 34. | Bgn | -1.09 | 0.0417 | Down |
| 35. | Tnfaip6 | -1.09 | 0.0007 | Down |
| 36. | Cpxm1 | -1.13 | 0.0006 | Down |
| 37. | Antxr1 | -1.14 | 0.0375 | Down |
| 38. | Aspn | -1.17 | 0.0604 | Down |
| 39. | Itm2a | -1.19 | 0.0003 | Down |
| 40. | Lgi2 | -1.19 | 0.0988 | Down |
| 41. | Fstl1 | -1.20 | 0.0170 | Down |
| 42. | Pcsk5 | -1.22 | 0.0708 | Down |
| 43. | Gpc3 | -1.23 | 0.0430 | Down |
| 44. | Rian | -1.32 | 0.0853 | Down |

| | | | | |
|---|---|---|---|---|
| 45. | Csgalnact1 | -1.38 | 0.0102 | Down |
| 46. | Adamts2 | -1.40 | 0.0648 | Down |
| 47. | Has1 | -1.46 | 0.0333 | Down |
| 48. | Gm13889 | -1.46 | 0.0908 | Down |
| 49. | Nrcam | -1.47 | 0.0535 | Down |
| 50. | Neto2 | -1.48 | 0.0975 | Down |
| 51. | Cspg4 | -1.49 | 0.0360 | Down |
| 52. | Adamts4 | -1.51 | 0.0162 | Down |
| 53. | Gm47283 | -1.54 | 0.0043 | Down |
| 54. | Gsto2 | -1.55 | 0.0329 | Down |
| 55. | Lama2 | -1.56 | 0.0097 | Down |
| 56. | Gm37069 | -1.57 | 0.0213 | Down |
| 57. | Lum | -1.57 | 0.0001 | Down |
| 58. | Pcdh17 | -1.57 | 0.0654 | Down |
| 59. | Gm16310 | -1.58 | 0.0648 | Down |
| 60. | Pgf | -1.61 | 0.0507 | Down |
| 61. | Abca8a | -1.63 | 0.0299 | Down |
| 62. | Trim69 | -1.74 | 0.0573 | Down |
| 63. | Stc1 | -1.77 | 0.0462 | Down |
| 64. | Scn7a | -1.77 | 0.0667 | Down |
| 65. | Ttc21a | -1.78 | 0.0125 | Down |
| 66. | Col23a1 | -1.79 | 0.0573 | Down |
| 67. | Adamtsl1 | -1.81 | 0.0880 | Down |
| 68. | Iqcn | -1.81 | 0.0000 | Down |
| 69. | Podn | -1.87 | 0.0375 | Down |
| 70. | Sema5a | -1.88 | 0.0853 | Down |
| 71. | Galnt16 | -1.89 | 0.0454 | Down |
| 72. | Gfra1 | -1.90 | 0.0581 | Down |
| 73. | Ncam1 | -2.01 | 0.0574 | Down |
| 74. | Raver2 | -2.03 | 0.0000 | Down |
| 75. | Gm48565 | -2.07 | 0.0160 | Down |
| 76. | Ntrk2 | -2.10 | 0.0006 | Down |
| 77. | Plp1 | -2.27 | 0.0550 | Down |
| 78. | Epha3 | -2.33 | 0.0048 | Down |
| 79. | Apod | -2.34 | 0.0000 | Down |
| 80. | Sprr2a3 | -2.39 | 0.0516 | Down |
| 81. | Cemip | -2.49 | 0.0003 | Down |
| 82. | Hpca | -2.53 | 0.0573 | Down |
| 83. | Bdh2 | -2.54 | 0.0091 | Down |
| 84. | Col28a1 | -2.60 | 0.0003 | Down |
| 85. | Adra2b | -2.64 | 0.0723 | Down |
| 86. | Gm33115 | -2.67 | 0.0788 | Down |
| 87. | Tacr1 | -3.01 | 0.0003 | Down |
| 88. | Slc6a17 | -3.06 | 0.0505 | Down |
| 89. | Cd207 | -3.81 | 0.0000 | Down |
| 90. | Gldn | -3.81 | 0.0122 | Down |
| 91. | Asxl3 | -4.68 | 0.0334 | Down |
| 92. | Gm49296 | -4.98 | 0.0648 | Down |

| | | | | |
|---|---|---|---|---|
| 93. | Cdh19 | -5.12 | 0.0007 | Down |

Supplementary table 2: Hallmark Gene Set Enrichment analysis (GSEA) at day fifteen post tumor challenge

| | Description | Set Size | Enrichment Score | NES | p value | p adjusted value | q value |
|---|---|---|---|---|---|---|---|
| 1 | HALLMARK_E2F_TARGETS | 199 | 0.569 | 2.274 | 1E-10 | 1.25E-09 | 6.0526E-10 |
| 2 | HALLMARK_OXIDATIVE_PHOSPHORYLATION | 185 | 0.575 | 2.273 | 1E-10 | 1.25E-09 | 6.0526E-10 |
| 3 | HALLMARK_MYC_TARGETS_V1 | 195 | 0.549 | 2.179 | 1E-10 | 1.25E-09 | 6.0526E-10 |
| 4 | HALLMARK_ALLOGRAFT_REJECTION | 194 | 0.494 | 1.959 | 2.5443E-08 | 2.1202E-07 | 1.0266E-07 |
| 5 | HALLMARK_DNA_REPAIR | 148 | 0.493 | 1.901 | 9.0686E-07 | 5.6679E-06 | 2.7444E-06 |
| 6 | HALLMARK_MYC_TARGETS_V2 | 57 | 0.569 | 1.885 | 0.00014413 | 0.00042391 | 0.00020526 |
| 7 | HALLMARK_FATTY_ACID_METABOLISM | 150 | 0.442 | 1.709 | 5.0514E-05 | 0.00016838 | 8.1532E-05 |
| 8 | HALLMARK_INTERFERON_ALPHA_RESPONSE | 98 | 0.417 | 1.511 | 0.00722897 | 0.01571514 | 0.00760944 |
| 9 | HALLMARK_G2M_CHECKPOINT | 197 | 0.371 | 1.482 | 0.00149944 | 0.0039459 | 0.00191065 |
| 10 | HALLMARK_INTERFERON_GAMMA_RESPONSE | 199 | 0.366 | 1.461 | 0.00226501 | 0.00539289 | 0.00261129 |
| 11 | HALLMARK_ADIPOGENESIS | 198 | 0.364 | 1.455 | 0.00291215 | 0.00661852 | 0.00320476 |
| 12 | HALLMARK_PEROXISOME | 96 | 0.392 | 1.415 | 0.02041402 | 0.03925774 | 0.01900901 |
| 13 | HALLMARK_P53_PATHWAY | 197 | 0.332 | 1.324 | 0.01398283 | 0.02796566 | 0.01354127 |
| 14 | HALLMARK_APOPTOSIS | 159 | -0.382 | -1.476 | 0.00785238 | 0.01635912 | 0.00792126 |
| 15 | HALLMARK_KRAS_SIGNALING_DN | 176 | -0.396 | -1.550 | 0.00172146 | 0.00430366 | 0.00208388 |
| 16 | HALLMARK_ESTROGEN_RESPONSE_EARLY | 190 | -0.402 | -1.585 | 0.00074581 | 0.0020717 | 0.00100314 |
| 17 | HALLMARK_APICAL_JUNCTION | 193 | -0.435 | -1.718 | 6.4462E-05 | 0.00020144 | 9.7541E-05 |
| 18 | HALLMARK_TNFA_SIGNALING_VIA_NFKB | 197 | -0.437 | -1.723 | 4.8185E-05 | 0.00016838 | 8.1532E-05 |
| 19 | HALLMARK_INFLAMMATORY_RESPONSE | 195 | -0.439 | -1.736 | 3.273E-05 | 0.00013638 | 6.6034E-05 |
| 20 | HALLMARK_HYPOXIA | 193 | -0.453 | -1.788 | 1.5079E-05 | 6.8542E-05 | 3.3189E-05 |
| 21 | HALLMARK_KRAS_SIGNALING_UP | 197 | -0.453 | -1.789 | 1.1039E-05 | 5.5195E-05 | 2.6726E-05 |
| 22 | HALLMARK_MYOGENESIS | 191 | -0.497 | -1.961 | 3.6721E-07 | 2.623E-06 | 1.2701E-06 |
| 23 | HALLMARK_COAGULATION | 130 | -0.529 | -1.994 | 1.0938E-06 | 6.0766E-06 | 2.9424E-06 |
| 24 | HALLMARK_ANGIOGENESIS | 35 | -0.719 | -2.132 | 4.0057E-05 | 0.00015406 | 7.4599E-05 |

Supplementary table 3: Top 15 gene ontology (GO) Gene Sets ranked by NES at day fifteen post tumor challenge

| | Description | Set Size | Enrichment Score | NES | p value | p adjusted value | q value |
|---|---|---|---|---|---|---|---|
| 1 | GOMF_STRUCTURAL_CONSTITUENT_OF_RIBOSOME | 153 | 0.760 | 2.952 | 1E-10 | 1.4221E-08 | 1.0516E-08 |
| 2 | GOCC_RIBOSOMAL_SUBUNIT | 177 | 0.736 | 2.905 | 1E-10 | 1.4221E-08 | 1.0516E-08 |
| 3 | GOCC_LARGE_RIBOSOMAL_SUBUNIT | 107 | 0.746 | 2.744 | 1E-10 | 1.4221E-08 | 1.0516E-08 |
| 4 | GOCC_CYTOSOLIC_LARGE_RIBOSOMAL_SUBUNIT | 51 | 0.839 | 2.735 | 1E-10 | 1.4221E-08 | 1.0516E-08 |
| 5 | GOBP_CYTOPLASMIC_TRANSLATION | 156 | 0.696 | 2.709 | 1E-10 | 1.4221E-08 | 1.0516E-08 |
| 6 | GOCC_MITOCHONDRIAL_PROTEIN_CONTAINING_COMPLEX | 260 | 0.635 | 2.634 | 1E-10 | 1.4221E-08 | 1.0516E-08 |
| 7 | GOCC_RIBOSOME | 222 | 0.633 | 2.578 | 1E-10 | 1.4221E-08 | 1.0516E-08 |
| 8 | GOCC_SMALL_RIBOSOMAL_SUBUNIT | 73 | 0.736 | 2.547 | 1E-10 | 1.4221E-08 | 1.0516E-08 |
| 9 | GOBP_MITOCHONDRIAL_GENE_EXPRESSION | 164 | 0.646 | 2.530 | 1E-10 | 1.4221E-08 | 1.0516E-08 |
| 10 | GOBP_MITOCHONDRIAL_TRANSLATION | 127 | 0.671 | 2.520 | 1E-10 | 1.4221E-08 | 1.0516E-08 |
| 11 | GOCC_CYTOSOLIC_RIBOSOME | 110 | 0.680 | 2.514 | 1E-10 | 1.4221E-08 | 1.0516E-08 |
| 12 | GOCC_T_CELL_RECEPTOR_COMPLEX | 80 | 0.708 | 2.485 | 1E-10 | 1.4221E-08 | 1.0516E-08 |
| 13 | GOCC_CYTOSOLIC_SMALL_RIBOSOMAL_SUBUNIT | 41 | 0.795 | 2.484 | 1E-10 | 1.4221E-08 | 1.0516E-08 |
| 14 | HP_RETICULOCYTOPENIA | 36 | 0.813 | 2.482 | 1E-10 | 1.4221E-08 | 1.0516E-08 |
| 15 | GOCC_MITOCHONDRIAL_MATRIX | 463 | 0.564 | 2.471 | 1E-10 | 1.4221E-08 | 1.0516E-08 |
| 16 | GOMF_EXTRACELLULAR_MATRIX_STRUCTURAL_CONSTITUENT | 141 | -0.760 | -2.889 | 1E-10 | 1.42213E-08 | 1.05165E-08 |
| 17 | GOCC_COLLAGEN_CONTAINING_EXTRACELLULAR_MATRIX | 387 | -0.673 | -2.884 | 1E-10 | 1.42213E-08 | 1.05165E-08 |
| 18 | GOBP_EXTERNAL_ENCAPSULATING_STRUCTURE_ORGANIZATION | 307 | -0.663 | -2.778 | 1E-10 | 1.42213E-08 | 1.05165E-08 |
| 19 | GOBP_COLLAGEN_FIBRIL_ORGANIZATION | 65 | -0.799 | -2.675 | 1E-10 | 1.42213E-08 | 1.05165E-08 |
| 20 | GOMF_GLYCOSAMINOGLYCAN_BINDING | 215 | -0.654 | -2.620 | 1E-10 | 1.42213E-08 | 1.05165E-08 |
| 21 | GOCC_COLLAGEN_TRIMER | 72 | -0.754 | -2.569 | 1E-10 | 1.42213E-08 | 1.05165E-08 |

| | | | | | | | |
|---|---|---|---|---|---|---|---|
| 22 | GOMF_COLLAGEN_BINDING | 66 | -0.755 | -2.529 | 1E-10 | 1.42213E-08 | 1.05165E-08 |
| 23 | GOCC_BASEMENT_MEMBRANE | 86 | -0.707 | -2.512 | 1E-10 | 1.42213E-08 | 1.05165E-08 |
| 24 | GOMF_HEPARIN_BINDING | 150 | -0.645 | -2.467 | 1E-10 | 1.42213E-08 | 1.05165E-08 |
| 25 | GOMF_EXTRACELLULAR_MATRIX_STRUCTURAL_CONSTITUENT_CONFERRING_TENSILE_STRENGTH | 39 | -0.787 | -2.425 | 3.20193E-08 | 2.27679E-06 | 1.68365E-06 |
| 26 | GOBP_CARTILAGE_DEVELOPMENT | 192 | -0.603 | -2.398 | 1E-10 | 1.42213E-08 | 1.05165E-08 |
| 27 | HP_ABNORMALITY_OF_HAND_JOINT_MOBILITY | 47 | -0.755 | -2.386 | 2.23148E-08 | 1.6644E-06 | 1.2308E-06 |
| 28 | GOBP_HOMOPHILIC_CELL_ADHESION_VIA_PLASMA_MEMBRANE_ADHESION_MOLECULES | 143 | -0.624 | -2.374 | 1E-10 | 1.42213E-08 | 1.05165E-08 |
| 29 | GOMF_SERINE_TYPE_ENDOPEPTIDASE_INHIBITOR_ACTIVITY | 71 | -0.694 | -2.372 | 1.09198E-08 | 9.17092E-07 | 6.78177E-07 |
| 30 | HP_ATROPHIC_SCARS | 36 | -0.790 | -2.357 | 1.70961E-07 | 9.16317E-06 | 6.77605E-06 |

Supplementary table 4: Top 10 KEGG pathways ranked by NES at day fifteen post tumor challenge

| No | ID | Description | Set Size | Enrichment Score | NES | p value | p adjusted value | q value |
|---|---|---|---|---|---|---|---|---|
| 1 | mmu03010 | Ribosome | 172 | 0.707 | 2.775 | 1E-10 | 6.28254E-09 | 3.89571E-09 |
| 2 | mmu05340 | Primary immunodeficiency | 34 | 0.723 | 2.184 | 3.82928E-06 | 6.92989E-05 | 4.29712E-05 |
| 3 | mmu03030 | DNA replication | 36 | 0.707 | 2.174 | 7.7273E-06 | 0.00012223 | 7.5795E-05 |
| 4 | mmu04612 | Antigen processing and presentation | 73 | 0.618 | 2.098 | 9.9353E-08 | 3.1432E-06 | 1.949E-06 |
| 5 | mmu05171 | Coronavirus disease - COVID-19 | 260 | 0.517 | 2.126 | 1E-10 | 6.2825E-09 | 3.8957E-09 |
| 6 | mmu04512 | ECM-receptor interaction | 84 | -0.685 | -2.420 | 1.0832E-10 | 6.2825E-09 | 3.8957E-09 |
| 7 | mmu04820 | Cytoskeleton in muscle cells | 219 | -0.593 | -2.402 | 1E-10 | 6.2825E-09 | 3.8957E-09 |
| 8 | mmu04974 | Protein digestion and absorption | 90 | -0.666 | -2.388 | 1.2473E-09 | 5.4257E-08 | 3.3644E-08 |
| 9 | mmu04510 | Focal adhesion | 196 | -0.547 | -2.186 | 2.2917E-10 | 1.1393E-08 | 7.0646E-09 |
| 10 | mmu04151 | PI3K-Akt signaling pathway | 337 | -0.502 | -2.116 | 1E-10 | 6.2825E-09 | 3.8957E-09 |

Supplementary table 5: Differentially expressed genes at day thirty post tumor innoculation

| | Gene name | log2FC | padj | regulation |
|---|---|---|---|---|
| 1. | Crisp3 | 7.70 | 0.0117 | Up |
| 2. | Fcer2a | 7.20 | 0.0148 | Up |
| 3. | Prlr | 6.78 | 0.0376 | Up |
| 4. | Foxn1 | 5.91 | 0.0424 | Up |
| 5. | Fcmr | 5.86 | 0.0126 | Up |
| 6. | Slc28a2b | 5.72 | 0.0969 | Up |
| 7. | Klk6 | 5.31 | 0.0032 | Up |
| 8. | Pax5 | 5.15 | 0.0323 | Up |
| 9. | Cd79a | 5.03 | 0.0030 | Up |
| 10. | Blk | 5.01 | 0.0062 | Up |
| 11. | Cd19 | 4.74 | 0.0054 | Up |
| 12. | Cacna1i | 4.65 | 0.0109 | Up |
| 13. | Srpk3 | 4.61 | 0.0137 | Up |
| 14. | Ms4a1 | 4.45 | 0.0147 | Up |
| 15. | Dapl1 | 4.44 | 0.0603 | Up |
| 16. | Cntfr | 4.37 | 0.0003 | Up |
| 17. | Samd5 | 4.30 | 0.0086 | Up |
| 18. | Slc6a19 | 4.29 | 0.0969 | Up |
| 19. | Lcn4 | 4.17 | 0.0441 | Up |
| 20. | Pkhd1 | 4.17 | 0.0148 | Up |
| 21. | Tdrp | 4.03 | 0.0177 | Up |
| 22. | Map3k21 | 3.99 | 0.0998 | Up |
| 23. | Acan | 3.97 | 0.0427 | Up |
| 24. | Fcrl1 | 3.89 | 0.0236 | Up |
| 25. | Pou2af1 | 3.87 | 0.0510 | Up |
| 26. | Lef1 | 3.86 | 0.0882 | Up |
| 27. | Chst3 | 3.72 | 0.0151 | Up |
| 28. | Wnt7a | 3.71 | 0.0003 | Up |
| 29. | Kcnh2 | 3.69 | 0.0410 | Up |
| 30. | Tslp | 3.63 | 0.0109 | Up |
| 31. | Tnfrsf13c | 3.52 | 0.0427 | Up |
| 32. | Nsg2 | 3.47 | 0.0522 | Up |
| 33. | Tbxa2r | 3.43 | 0.0522 | Up |
| 34. | Rapgef4 | 3.42 | 0.0137 | Up |
| 35. | Scd1 | 3.29 | 0.0026 | Up |
| 36. | Vipr1 | 3.24 | 0.0758 | Up |
| 37. | Fcrla | 3.18 | 0.0439 | Up |
| 38. | Dok7 | 3.16 | 0.0802 | Up |
| 39. | Tmie | 3.16 | 0.0127 | Up |
| 40. | H2-Ob | 3.09 | 0.0435 | Up |
| 41. | St8sia6 | 3.06 | 0.0198 | Up |
| 42. | Pik3ip1 | 3.00 | 0.0117 | Up |
| 43. | Ccbe1 | 2.95 | 0.0174 | Up |
| 44. | Scml4 | 2.94 | 0.0557 | Up |

| | | | | |
|---|---|---|---|---|
| 45. | S1pr1 | 2.90 | 0.0356 | Up |
| 46. | Cmah | 2.89 | 0.0427 | Up |
| 47. | Pik3c2b | 2.86 | 0.0054 | Up |
| 48. | Cldn6 | 2.84 | 0.0687 | Up |
| 49. | Ebf1 | 2.82 | 0.0142 | Up |
| 50. | Cxcr5 | 2.75 | 0.0622 | Up |
| 51. | Garnl3 | 2.74 | 0.0435 | Up |
| 52. | Dnah8 | 2.71 | 0.0896 | Up |
| 53. | Il5ra | 2.65 | 0.0244 | Up |
| 54. | Nudt6 | 2.59 | 0.0736 | Up |
| 55. | Tcf7 | 2.55 | 0.0712 | Up |
| 56. | Cd79b | 2.53 | 0.0765 | Up |
| 57. | Sema3c | 2.52 | 0.0743 | Up |
| 58. | Hoxb9 | 2.50 | 0.0342 | Up |
| 59. | Rhpn2 | 2.50 | 0.0248 | Up |
| 60. | Cd22 | 2.44 | 0.0882 | Up |
| 61. | Ankrd1 | 2.42 | 0.0196 | Up |
| 62. | Sfrp1 | 2.37 | 0.0806 | Up |
| 63. | Dzip1 | 2.34 | 0.0874 | Up |
| 64. | Bach2 | 2.34 | 0.0497 | Up |
| 65. | Nkx2-3 | 2.34 | 0.0356 | Up |
| 66. | Sbk1 | 2.33 | 0.0117 | Up |
| 67. | Rasgrp3 | 2.29 | 0.0557 | Up |
| 68. | Palm3 | 2.25 | 0.0904 | Up |
| 69. | Als2cl | 2.24 | 0.0057 | Up |
| 70. | Arhgef18 | 2.21 | 0.0148 | Up |
| 71. | Patj | 2.19 | 0.0378 | Up |
| 72. | Itm2a | 2.18 | 0.0141 | Up |
| 73. | Ltbp2 | 2.18 | 0.0955 | Up |
| 74. | Bcl7a | 2.17 | 0.0347 | Up |
| 75. | Syde2 | 2.15 | 0.0344 | Up |
| 76. | Acer2 | 2.13 | 0.0522 | Up |
| 77. | Ift80 | 2.12 | 0.0439 | Up |
| 78. | Card6 | 2.11 | 0.0542 | Up |
| 79. | Snn | 2.09 | 0.0802 | Up |
| 80. | Sorbs2 | 2.08 | 0.0682 | Up |
| 81. | Pitpnm2 | 2.08 | 0.0448 | Up |
| 82. | Zfp354c | 2.06 | 0.0944 | Up |
| 83. | Dus4l | 2.03 | 0.0614 | Up |
| 84. | Ralgps2 | 1.97 | 0.0381 | Up |
| 85. | Elovl7 | 1.96 | 0.0356 | Up |
| 86. | Cd55 | 1.95 | 0.0255 | Up |
| 87. | Muc1 | 1.90 | 0.0234 | Up |
| 88. | Chrnb1 | 1.89 | 0.0694 | Up |
| 89. | Rhobtb2 | 1.86 | 0.0834 | Up |
| 90. | Fgf13 | 1.84 | 0.0322 | Up |
| 91. | Lrig1 | 1.83 | 0.0373 | Up |
| 92. | Ppp2r2c | 1.82 | 0.0751 | Up |

| | | | | |
|---|---|---|---|---|
| 93. | Tgfbr3 | 1.78 | 0.0407 | Up |
| 94. | Kifc2 | 1.78 | 0.0462 | Up |
| 95. | Nckap5l | 1.76 | 0.0396 | Up |
| 96. | Adam12 | 1.75 | 0.0885 | Up |
| 97. | Slc35f2 | 1.73 | 0.0660 | Up |
| 98. | Ankrd26 | 1.69 | 0.0356 | Up |
| 99. | Hid1 | 1.66 | 0.0284 | Up |
| 100. | Angel1 | 1.64 | 0.0372 | Up |
| 101. | Cyb5rl | 1.60 | 0.0435 | Up |
| 102. | Ephb6 | 1.60 | 0.0692 | Up |
| 103. | Stx1a | 1.58 | 0.0943 | Up |
| 104. | Rras2 | 1.57 | 0.0395 | Up |
| 105. | Pgap1 | 1.55 | 0.0708 | Up |
| 106. | Pxylp1 | 1.54 | 0.0202 | Up |
| 107. | Mapk11 | 1.47 | 0.0203 | Up |
| 108. | Slc12a2 | 1.47 | 0.0320 | Up |
| 109. | Znrf3 | 1.45 | 0.0740 | Up |
| 110. | Tnfrsf22 | 1.45 | 0.0557 | Up |
| 111. | Rab3ip | 1.43 | 0.0925 | Up |
| 112. | Auts2 | 1.43 | 0.0392 | Up |
| 113. | Lmntd2 | 1.43 | 0.0699 | Up |
| 114. | Tnfrsf10b | 1.42 | 0.0557 | Up |
| 115. | Enpp1 | 1.38 | 0.0407 | Up |
| 116. | Fitm2 | 1.38 | 0.0834 | Up |
| 117. | Strbp | 1.37 | 0.0630 | Up |
| 118. | L2hgdh | 1.36 | 0.0508 | Up |
| 119. | Csrnp2 | 1.35 | 0.0943 | Up |
| 120. | Tarbp1 | 1.35 | 0.0427 | Up |
| 121. | Mettl8 | 1.34 | 0.0890 | Up |
| 122. | Actn1 | 1.33 | 0.0041 | Up |
| 123. | Cep68 | 1.33 | 0.0086 | Up |
| 124. | Adgrl1 | 1.32 | 0.0026 | Up |
| 125. | Acad10 | 1.31 | 0.0959 | Up |
| 126. | Miga1 | 1.28 | 0.0091 | Up |
| 127. | Tecpr1 | 1.27 | 0.0712 | Up |
| 128. | Prickle1 | 1.26 | 0.0691 | Up |
| 129. | Bbs9 | 1.22 | 0.0433 | Up |
| 130. | Rcbtb1 | 1.22 | 0.0834 | Up |
| 131. | Kdsr | 1.20 | 0.0362 | Up |
| 132. | Cdh24 | 1.20 | 0.0350 | Up |
| 133. | Fam120c | 1.18 | 0.0090 | Up |
| 134. | Pfn2 | 1.18 | 0.0356 | Up |
| 135. | Eno3 | 1.16 | 0.0435 | Up |
| 136. | Golm2 | 1.14 | 0.0422 | Up |
| 137. | Pkd1 | 1.11 | 0.0126 | Up |
| 138. | Heg1 | 1.11 | 0.0806 | Up |
| 139. | Hdac7 | 1.10 | 0.0954 | Up |
| 140. | Mccc2 | 1.09 | 0.0955 | Up |

| | | | | |
|---|---|---|---|---|
| 141. | Utrn | 1.09 | 0.0830 | Up |
| 142. | Zfp827 | 1.09 | 0.0148 | Up |
| 143. | Grwd1 | 1.06 | 0.0768 | Up |
| 144. | Msh2 | 1.05 | 0.0439 | Up |
| 145. | Ppif | 1.05 | 0.0431 | Up |
| 146. | Qtrt2 | 1.04 | 0.0441 | Up |
| 147. | Ttc13 | 1.04 | 0.0242 | Up |
| 148. | Stambpl1 | 1.03 | 0.0198 | Up |
| 149. | Cdr2 | 1.03 | 0.0048 | Up |
| 150. | Ttc3 | 1.03 | 0.0323 | Up |
| 151. | Sptbn1 | 1.02 | 0.0054 | Up |
| 152. | Pus7 | 1.01 | 0.0630 | Up |
| 153. | Chpf | 1.01 | 0.0117 | Up |
| 154. | Zfp12 | 1.00 | 0.0522 | Up |
| 155. | Ankrd10 | 0.99 | 0.0467 | Up |
| 156. | Ttc27 | 0.98 | 0.0522 | Up |
| 157. | Rcan3 | 0.98 | 0.0164 | Up |
| 158. | Tpk1 | 0.98 | 0.0896 | Up |
| 159. | Adcy3 | 0.98 | 0.0086 | Up |
| 160. | Slc5a6 | 0.97 | 0.0925 | Up |
| 161. | Rab11fip3 | 0.96 | 0.0466 | Up |
| 162. | Gnl3 | 0.96 | 0.0892 | Up |
| 163. | Psip1 | 0.96 | 0.0806 | Up |
| 164. | Klhl42 | 0.96 | 0.0522 | Up |
| 165. | Dhodh | 0.95 | 0.0483 | Up |
| 166. | Nob1 | 0.93 | 0.0699 | Up |
| 167. | Itfg2 | 0.92 | 0.0536 | Up |
| 168. | Wdr74 | 0.92 | 0.0686 | Up |
| 169. | Ftsj3 | 0.91 | 0.0469 | Up |
| 170. | Slc7a6 | 0.91 | 0.0048 | Up |
| 171. | Utp20 | 0.89 | 0.0959 | Up |
| 172. | Pabpc4 | 0.87 | 0.0237 | Up |
| 173. | Tshz1 | 0.85 | 0.0376 | Up |
| 174. | Atad3a | 0.85 | 0.0496 | Up |
| 175. | Polr1a | 0.84 | 0.0834 | Up |
| 176. | Tmem245 | 0.84 | 0.0237 | Up |
| 177. | Trmt1 | 0.83 | 0.0331 | Up |
| 178. | Cplane1 | 0.83 | 0.0806 | Up |
| 179. | Tbcel | 0.83 | 0.0806 | Up |
| 180. | Pwp2 | 0.82 | 0.0698 | Up |
| 181. | Abhd2 | 0.82 | 0.0237 | Up |
| 182. | Zdhhc8 | 0.80 | 0.0091 | Up |
| 183. | Dhx33 | 0.80 | 0.0312 | Up |
| 184. | Aasdhppt | 0.77 | 0.0943 | Up |
| 185. | Zbtb4 | 0.76 | 0.0031 | Up |
| 186. | Fam210a | 0.73 | 0.0806 | Up |
| 187. | Mtfr1 | 0.73 | 0.0594 | Up |
| 188. | Zfp664 | 0.73 | 0.0149 | Up |

| | | | | |
|---|---|---|---|---|
| 189. | Fam199x | 0.73 | 0.0378 | Up |
| 190. | Esyt2 | 0.72 | 0.0692 | Up |
| 191. | Atic | 0.72 | 0.0516 | Up |
| 192. | Galnt7 | 0.71 | 0.0958 | Up |
| 193. | Mbtps1 | 0.71 | 0.0198 | Up |
| 194. | Uba52 | 0.69 | 0.0539 | Up |
| 195. | Iars2 | 0.69 | 0.0884 | Up |
| 196. | Cad | 0.68 | 0.0244 | Up |
| 197. | Gid4 | 0.68 | 0.0331 | Up |
| 198. | D030056L22Rik | 0.67 | 0.0428 | Up |
| 199. | Gnpat | 0.63 | 0.0522 | Up |
| 200. | Ubqln2 | 0.63 | 0.0122 | Up |
| 201. | Lrpprc | 0.62 | 0.0557 | Up |
| 202. | Pygo2 | 0.62 | 0.0407 | Up |
| 203. | Prepl | 0.62 | 0.0395 | Up |
| 204. | Men1 | 0.60 | 0.0435 | Up |
| 205. | Fam98a | 0.59 | 0.0806 | Up |
| 206. | Ippk | 0.58 | 0.0567 | Up |
| 207. | Akap11 | 0.57 | 0.0813 | Up |
| 208. | Cers5 | 0.53 | 0.0496 | Up |
| 209. | Rhog | -0.51 | 0.0828 | Down |
| 210. | Ube2e3 | -0.52 | 0.0678 | Down |
| 211. | Tnfaip8 | -0.53 | 0.0885 | Down |
| 212. | Shkbp1 | -0.53 | 0.0943 | Down |
| 213. | H3f3b | -0.54 | 0.0497 | Down |
| 214. | Mrpl52 | -0.54 | 0.0465 | Down |
| 215. | Psme1 | -0.55 | 0.0896 | Down |
| 216. | Gna13 | -0.55 | 0.0582 | Down |
| 217. | Tet2 | -0.56 | 0.0952 | Down |
| 218. | Tbc1d23 | -0.56 | 0.0997 | Down |
| 219. | Cox6b1 | -0.57 | 0.0826 | Down |
| 220. | Rtf2 | -0.59 | 0.0137 | Down |
| 221. | Traf3 | -0.59 | 0.0367 | Down |
| 222. | Ccdc12 | -0.60 | 0.0896 | Down |
| 223. | Capzb | -0.60 | 0.0435 | Down |
| 224. | Chmp2a | -0.62 | 0.0441 | Down |
| 225. | Actb | -0.62 | 0.0091 | Down |
| 226. | Tank | -0.62 | 0.0250 | Down |
| 227. | Sem1 | -0.62 | 0.0350 | Down |
| 228. | Slc25a20 | -0.63 | 0.0999 | Down |
| 229. | Psme2 | -0.64 | 0.0749 | Down |
| 230. | Zfp263 | -0.65 | 0.0438 | Down |
| 231. | Erp29 | -0.67 | 0.0462 | Down |
| 232. | Zcchc2 | -0.67 | 0.0889 | Down |
| 233. | Psmb8 | -0.68 | 0.0081 | Down |
| 234. | Itm2b | -0.68 | 0.0806 | Down |
| 235. | Rbbp8 | -0.69 | 0.0356 | Down |
| 236. | Pstpip1 | -0.70 | 0.0639 | Down |

| | | | | |
|---|---|---|---|---|
| 237. | H2-T23 | -0.70 | 0.0142 | Down |
| 238. | Chmp4b | -0.70 | 0.0030 | Down |
| 239. | Dusp16 | -0.71 | 0.0356 | Down |
| 240. | Ptpn1 | -0.71 | 0.0068 | Down |
| 241. | Arap1 | -0.72 | 0.0411 | Down |
| 242. | Gnpda1 | -0.72 | 0.0897 | Down |
| 243. | Myo1g | -0.73 | 0.0670 | Down |
| 244. | H2-DMb1 | -0.74 | 0.0983 | Down |
| 245. | Ap2s1 | -0.74 | 0.0959 | Down |
| 246. | Tmem170b | -0.75 | 0.0127 | Down |
| 247. | Fcho2 | -0.75 | 0.0676 | Down |
| 248. | Cep85l | -0.75 | 0.0148 | Down |
| 249. | Snx20 | -0.75 | 0.0198 | Down |
| 250. | Ric1 | -0.77 | 0.0411 | Down |
| 251. | Casp6 | -0.77 | 0.0850 | Down |
| 252. | P2rx4 | -0.78 | 0.0089 | Down |
| 253. | Mcl1 | -0.79 | 0.0688 | Down |
| 254. | Kctd12 | -0.80 | 0.0196 | Down |
| 255. | Gsdmd | -0.80 | 0.0809 | Down |
| 256. | Lsm10 | -0.80 | 0.0457 | Down |
| 257. | Plcg2 | -0.81 | 0.0015 | Down |
| 258. | Maf | -0.81 | 0.0670 | Down |
| 259. | Hpcal1 | -0.81 | 0.0013 | Down |
| 260. | Lcp2 | -0.81 | 0.0483 | Down |
| 261. | Ifi35 | -0.81 | 0.0951 | Down |
| 262. | Tnip2 | -0.82 | 0.0604 | Down |
| 263. | Lgals9 | -0.83 | 0.0196 | Down |
| 264. | Dapp1 | -0.83 | 0.0985 | Down |
| 265. | Smagp | -0.84 | 0.0557 | Down |
| 266. | Rnf34 | -0.84 | 0.0347 | Down |
| 267. | Cd52 | -0.85 | 0.0402 | Down |
| 268. | Tor1aip1 | -0.85 | 0.0255 | Down |
| 269. | Rab8b | -0.87 | 0.0711 | Down |
| 270. | Hmgcl | -0.87 | 0.0930 | Down |
| 271. | Irak2 | -0.88 | 0.0428 | Down |
| 272. | Slc35f6 | -0.88 | 0.0603 | Down |
| 273. | Taldo1 | -0.88 | 0.0913 | Down |
| 274. | Myd88 | -0.89 | 0.0299 | Down |
| 275. | Tap1 | -0.89 | 0.0039 | Down |
| 276. | Lpxn | -0.89 | 0.0191 | Down |
| 277. | Il10rb | -0.89 | 0.0806 | Down |
| 278. | Tent5a | -0.90 | 0.0204 | Down |
| 279. | Syk | -0.91 | 0.0234 | Down |
| 280. | Prkcd | -0.92 | 0.0951 | Down |
| 281. | Napsa | -0.92 | 0.0518 | Down |
| 282. | Trim25 | -0.92 | 0.0435 | Down |
| 283. | Ovca2 | -0.93 | 0.0120 | Down |
| 284. | Ifnar2 | -0.94 | 0.0691 | Down |

| | | | | |
|---|---|---|---|---|
| 285. | Etv3 | -0.95 | 0.0676 | Down |
| 286. | Dbi | -0.95 | 0.0424 | Down |
| 287. | Ogfrl1 | -0.95 | 0.0603 | Down |
| 288. | Rgs1 | -0.95 | 0.0356 | Down |
| 289. | Pcyt1a | -0.95 | 0.0682 | Down |
| 290. | Lipe | -0.96 | 0.0237 | Down |
| 291. | Psmb9 | -0.97 | 0.0142 | Down |
| 292. | Parp10 | -0.98 | 0.0142 | Down |
| 293. | Cyth4 | -0.99 | 0.0237 | Down |
| 294. | Picalm | -1.00 | 0.0448 | Down |
| 295. | Apaf1 | -1.00 | 0.0408 | Down |
| 296. | Coro2a | -1.01 | 0.0323 | Down |
| 297. | Kif19a | -1.01 | 0.0682 | Down |
| 298. | Alas1 | -1.01 | 0.0885 | Down |
| 299. | Arhgap18 | -1.02 | 0.0490 | Down |
| 300. | Lmnb1 | -1.02 | 0.0682 | Down |
| 301. | Relt | -1.02 | 0.0676 | Down |
| 302. | Vav3 | -1.02 | 0.0408 | Down |
| 303. | Acot11 | -1.04 | 0.0884 | Down |
| 304. | Mdm2 | -1.04 | 0.0662 | Down |
| 305. | Rilpl2 | -1.04 | 0.0345 | Down |
| 306. | Nampt | -1.05 | 0.0542 | Down |
| 307. | Ticam1 | -1.05 | 0.0323 | Down |
| 308. | Plekho1 | -1.06 | 0.0493 | Down |
| 309. | Glipr2 | -1.06 | 0.0248 | Down |
| 310. | Slfn2 | -1.06 | 0.0723 | Down |
| 311. | B4galt5 | -1.08 | 0.0983 | Down |
| 312. | Id2 | -1.08 | 0.0237 | Down |
| 313. | Tasl | -1.08 | 0.0344 | Down |
| 314. | Ago4 | -1.09 | 0.0279 | Down |
| 315. | Nuak2 | -1.10 | 0.0511 | Down |
| 316. | Myo1f | -1.10 | 0.0768 | Down |
| 317. | Ncf4 | -1.10 | 0.0177 | Down |
| 318. | Adora2a | -1.11 | 0.0127 | Down |
| 319. | Zyx | -1.11 | 0.0592 | Down |
| 320. | P2ry14 | -1.12 | 0.0017 | Down |
| 321. | Synj1 | -1.12 | 0.0356 | Down |
| 322. | St3gal4 | -1.13 | 0.0395 | Down |
| 323. | Gbp6 | -1.13 | 0.0884 | Down |
| 324. | Gk | -1.13 | 0.0536 | Down |
| 325. | Marchf1 | -1.13 | 0.0373 | Down |
| 326. | Rin3 | -1.14 | 0.0367 | Down |
| 327. | Plekho2 | -1.14 | 0.0247 | Down |
| 328. | Cd200r1 | -1.14 | 0.0704 | Down |
| 329. | Rap1gap2 | -1.14 | 0.0925 | Down |
| 330. | Serpina3g | -1.15 | 0.0440 | Down |
| 331. | Fem1c | -1.15 | 0.0147 | Down |
| 332. | Nfil3 | -1.15 | 0.0414 | Down |

| | | | | |
|---|---|---|---|---|
| 333. | Gpr160 | -1.16 | 0.0622 | Down |
| 334. | Nfkbie | -1.16 | 0.0127 | Down |
| 335. | Etv6 | -1.17 | 0.0510 | Down |
| 336. | Rnf135 | -1.17 | 0.0992 | Down |
| 337. | Gpr65 | -1.17 | 0.0127 | Down |
| 338. | Casp4 | -1.17 | 0.0343 | Down |
| 339. | Bach1 | -1.17 | 0.0356 | Down |
| 340. | Fgr | -1.18 | 0.0356 | Down |
| 341. | Slc18a2 | -1.18 | 0.0236 | Down |
| 342. | Cyp27a1 | -1.18 | 0.0994 | Down |
| 343. | Tbx21 | -1.18 | 0.0768 | Down |
| 344. | Mxd1 | -1.19 | 0.0031 | Down |
| 345. | Rab3il1 | -1.21 | 0.0884 | Down |
| 346. | Spi1 | -1.21 | 0.0602 | Down |
| 347. | Ifi47 | -1.21 | 0.0433 | Down |
| 348. | Serpinb9 | -1.23 | 0.0059 | Down |
| 349. | Gbp3 | -1.23 | 0.0625 | Down |
| 350. | Il2ra | -1.25 | 0.0117 | Down |
| 351. | Ahr | -1.26 | 0.0142 | Down |
| 352. | Il18bp | -1.26 | 0.0441 | Down |
| 353. | Plekhg1 | -1.27 | 0.0994 | Down |
| 354. | Gpr55 | -1.27 | 0.0953 | Down |
| 355. | Tiam1 | -1.27 | 0.0148 | Down |
| 356. | Cited4 | -1.27 | 0.0765 | Down |
| 357. | Inka1 | -1.27 | 0.0407 | Down |
| 358. | Tbc1d8 | -1.28 | 0.0091 | Down |
| 359. | Arid3a | -1.28 | 0.0356 | Down |
| 360. | Ppfibp2 | -1.28 | 0.0924 | Down |
| 361. | Zfp366 | -1.28 | 0.0427 | Down |
| 362. | Dnaaf3 | -1.28 | 0.0894 | Down |
| 363. | Ncf2 | -1.30 | 0.0438 | Down |
| 364. | Cbfa2t3 | -1.31 | 0.0539 | Down |
| 365. | Ebi3 | -1.31 | 0.0424 | Down |
| 366. | Irf8 | -1.32 | 0.0086 | Down |
| 367. | Ube2l6 | -1.32 | 0.0667 | Down |
| 368. | Cndp2 | -1.32 | 0.0244 | Down |
| 369. | Oasl2 | -1.32 | 0.0969 | Down |
| 370. | Slc8b1 | -1.32 | 0.0924 | Down |
| 371. | Prf1 | -1.33 | 0.0407 | Down |
| 372. | Cd300ld | -1.34 | 0.0538 | Down |
| 373. | Tmem273 | -1.34 | 0.0983 | Down |
| 374. | Cd274 | -1.35 | 0.0407 | Down |
| 375. | Evi2a | -1.35 | 0.0042 | Down |
| 376. | Parp14 | -1.35 | 0.0467 | Down |
| 377. | Klre1 | -1.36 | 0.0486 | Down |
| 378. | Klra7 | -1.36 | 0.0427 | Down |
| 379. | Irf5 | -1.36 | 0.0127 | Down |
| 380. | Trafd1 | -1.37 | 0.0086 | Down |

| | | | | |
|---|---|---|---|---|
| 381. | Samhd1 | -1.37 | 0.0039 | Down |
| 382. | Ptgir | -1.38 | 0.0574 | Down |
| 383. | B4galt6 | -1.38 | 0.0959 | Down |
| 384. | Ifih1 | -1.39 | 0.0557 | Down |
| 385. | Irgm1 | -1.39 | 0.0331 | Down |
| 386. | Dhrs3 | -1.39 | 0.0435 | Down |
| 387. | Il10ra | -1.40 | 0.0310 | Down |
| 388. | Il15ra | -1.40 | 0.0356 | Down |
| 389. | Cysltr2 | -1.40 | 0.0286 | Down |
| 390. | Rasa4 | -1.40 | 0.0998 | Down |
| 391. | Nrros | -1.40 | 0.0062 | Down |
| 392. | Ifng | -1.41 | 0.0427 | Down |
| 393. | Casp1 | -1.41 | 0.0032 | Down |
| 394. | Chpt1 | -1.42 | 0.0198 | Down |
| 395. | Tifa | -1.42 | 0.0946 | Down |
| 396. | Metrnl | -1.43 | 0.0889 | Down |
| 397. | Ret | -1.44 | 0.0806 | Down |
| 398. | Pstpip2 | -1.44 | 0.0860 | Down |
| 399. | Fabp5 | -1.45 | 0.0198 | Down |
| 400. | C130050O18Rik | -1.45 | 0.0959 | Down |
| 401. | Oas3 | -1.46 | 0.0664 | Down |
| 402. | Fuom | -1.47 | 0.0994 | Down |
| 403. | Stat2 | -1.47 | 0.0557 | Down |
| 404. | Klri2 | -1.48 | 0.0089 | Down |
| 405. | S100a4 | -1.49 | 0.0924 | Down |
| 406. | Lst1 | -1.49 | 0.0474 | Down |
| 407. | Plac8 | -1.50 | 0.0117 | Down |
| 408. | Zfp971 | -1.51 | 0.0862 | Down |
| 409. | Irf7 | -1.51 | 0.0826 | Down |
| 410. | Cd40 | -1.52 | 0.0448 | Down |
| 411. | Ccr2 | -1.52 | 0.0407 | Down |
| 412. | Klrk1 | -1.52 | 0.0015 | Down |
| 413. | I830077J02Rik | -1.52 | 0.0142 | Down |
| 414. | Il15 | -1.52 | 0.0896 | Down |
| 415. | Wnt11 | -1.53 | 0.0806 | Down |
| 416. | Atp8b5 | -1.53 | 0.0312 | Down |
| 417. | Pirb | -1.53 | 0.0435 | Down |
| 418. | Rtp4 | -1.53 | 0.0989 | Down |
| 419. | Zbp1 | -1.54 | 0.0356 | Down |
| 420. | Apobec1 | -1.54 | 0.0323 | Down |
| 421. | Rnase6 | -1.54 | 0.0329 | Down |
| 422. | Clec4a3 | -1.55 | 0.0427 | Down |
| 423. | Naip6 | -1.55 | 0.0408 | Down |
| 424. | Tnfrsf8 | -1.55 | 0.0086 | Down |
| 425. | Zbtb32 | -1.55 | 0.0840 | Down |
| 426. | Klrb1c | -1.56 | 0.0254 | Down |
| 427. | Mafb | -1.57 | 0.0999 | Down |
| 428. | Mmp25 | -1.58 | 0.0344 | Down |

| | | | | |
|---|---|---|---|---|
| 429. | Cd244a | -1.58 | 0.0030 | Down |
| 430. | Xaf1 | -1.58 | 0.0330 | Down |
| 431. | Sowahc | -1.58 | 0.0362 | Down |
| 432. | Oas1a | -1.59 | 0.0557 | Down |
| 433. | Asb2 | -1.59 | 0.0445 | Down |
| 434. | Htr7 | -1.60 | 0.0778 | Down |
| 435. | Cd300a | -1.60 | 0.0137 | Down |
| 436. | Hck | -1.60 | 0.0126 | Down |
| 437. | Themis2 | -1.61 | 0.0004 | Down |
| 438. | Ifit1 | -1.62 | 0.0885 | Down |
| 439. | Fgl2 | -1.63 | 0.0625 | Down |
| 440. | Adora3 | -1.63 | 0.0524 | Down |
| 441. | Ccr5 | -1.64 | 0.0248 | Down |
| 442. | Lrrc25 | -1.64 | 0.0483 | Down |
| 443. | Ticam2 | -1.65 | 0.0141 | Down |
| 444. | Bnip5 | -1.65 | 0.0411 | Down |
| 445. | Gm5150 | -1.66 | 0.0356 | Down |
| 446. | Naip5 | -1.66 | 0.0039 | Down |
| 447. | Lilra6 | -1.67 | 0.0963 | Down |
| 448. | Ptpro | -1.68 | 0.0691 | Down |
| 449. | Phf11b | -1.68 | 0.0685 | Down |
| 450. | Ifi204 | -1.69 | 0.0720 | Down |
| 451. | Reps2 | -1.69 | 0.0946 | Down |
| 452. | Slfn5 | -1.71 | 0.0372 | Down |
| 453. | Rasgrp4 | -1.71 | 0.0163 | Down |
| 454. | Tmprss6 | -1.71 | 0.0924 | Down |
| 455. | Ly6c2 | -1.71 | 0.0356 | Down |
| 456. | Ovol2 | -1.72 | 0.0467 | Down |
| 457. | Septin3 | -1.73 | 0.0019 | Down |
| 458. | Adora2b | -1.73 | 0.0924 | Down |
| 459. | Oas1g | -1.73 | 0.0959 | Down |
| 460. | Fgd4 | -1.74 | 0.0203 | Down |
| 461. | Ly96 | -1.74 | 0.0005 | Down |
| 462. | Prr5l | -1.76 | 0.0198 | Down |
| 463. | Rgl1 | -1.77 | 0.0081 | Down |
| 464. | Ms4a4c | -1.77 | 0.0969 | Down |
| 465. | Cd300lb | -1.77 | 0.0356 | Down |
| 466. | Usp18 | -1.77 | 0.0462 | Down |
| 467. | Dgkg | -1.78 | 0.0759 | Down |
| 468. | Tlr9 | -1.78 | 0.0751 | Down |
| 469. | Rgs18 | -1.79 | 0.0306 | Down |
| 470. | Ifi211 | -1.79 | 0.0582 | Down |
| 471. | Pdcd1lg2 | -1.81 | 0.0137 | Down |
| 472. | Ssc4d | -1.81 | 0.0564 | Down |
| 473. | Rassf4 | -1.82 | 0.0039 | Down |
| 474. | Dhx58 | -1.82 | 0.0272 | Down |
| 475. | Gata2 | -1.83 | 0.0660 | Down |
| 476. | Mag | -1.84 | 0.0695 | Down |

| | | | | |
|---|---|---|---|---|
| 477. | Cmpk2 | -1.84 | 0.0172 | Down |
| 478. | Gm12185 | -1.85 | 0.0362 | Down |
| 479. | Oas2 | -1.86 | 0.0536 | Down |
| 480. | Ifi207 | -1.87 | 0.0427 | Down |
| 481. | Rsad2 | -1.89 | 0.0323 | Down |
| 482. | Slamf7 | -1.90 | 0.0126 | Down |
| 483. | Pira1 | -1.91 | 0.0395 | Down |
| 484. | Gbp5 | -1.91 | 0.0135 | Down |
| 485. | Isg15 | -1.92 | 0.0255 | Down |
| 486. | Klrg1 | -1.92 | 0.0042 | Down |
| 487. | Oasl1 | -1.95 | 0.0279 | Down |
| 488. | Fcgr1 | -1.95 | 0.0557 | Down |
| 489. | Cd209a | -1.95 | 0.0126 | Down |
| 490. | Mcpt2 | -1.96 | 0.0738 | Down |
| 491. | Nhsl2 | -1.97 | 0.0462 | Down |
| 492. | Phf11d | -1.98 | 0.0706 | Down |
| 493. | Klra9 | -1.98 | 0.0001 | Down |
| 494. | Gzme | -1.99 | 0.0466 | Down |
| 495. | Aldh1a2 | -1.99 | 0.0809 | Down |
| 496. | Phf11a | -2.00 | 0.0350 | Down |
| 497. | Ifit3b | -2.01 | 0.0317 | Down |
| 498. | Ms4a2 | -2.05 | 0.0427 | Down |
| 499. | Adgre4 | -2.05 | 0.0157 | Down |
| 500. | Calhm6 | -2.06 | 0.0885 | Down |
| 501. | Cxcl10 | -2.09 | 0.0148 | Down |
| 502. | Gpr141 | -2.15 | 0.0813 | Down |
| 503. | Siglecf | -2.15 | 0.0574 | Down |
| 504. | Tpsb2 | -2.15 | 0.0244 | Down |
| 505. | Mx2 | -2.17 | 0.0148 | Down |
| 506. | Bcl2l14 | -2.18 | 0.0259 | Down |
| 507. | Ifit3 | -2.20 | 0.0685 | Down |
| 508. | Gzmb | -2.27 | 0.0001 | Down |
| 509. | Cma1 | -2.28 | 0.0148 | Down |
| 510. | Mrgprb1 | -2.32 | 0.0121 | Down |
| 511. | Fcer1a | -2.33 | 0.0082 | Down |
| 512. | Ncr1 | -2.35 | 0.0258 | Down |
| 513. | F5 | -2.35 | 0.0148 | Down |
| 514. | Gm5431 | -2.36 | 0.0800 | Down |
| 515. | Gapt | -2.36 | 0.0015 | Down |
| 516. | Cpa3 | -2.37 | 0.0048 | Down |
| 517. | P2rx1 | -2.40 | 0.0780 | Down |
| 518. | Ccr3 | -2.43 | 0.0428 | Down |
| 519. | Mcpt4 | -2.46 | 0.0010 | Down |
| 520. | Retnlg | -2.47 | 0.0798 | Down |
| 521. | Cd209c | -2.48 | 0.0031 | Down |
| 522. | Cd200r3 | -2.50 | 0.0001 | Down |
| 523. | Ifit2 | -2.52 | 0.0237 | Down |
| 524. | Gzmc | -2.52 | 0.0001 | Down |

| | | | | |
|---|---|---|---|---|
| 525. | Pla2g3 | -2.53 | 0.0108 | Down |
| 526. | Clec4b1 | -2.56 | 0.0802 | Down |
| 527. | Clnk | -2.59 | 0.0439 | Down |
| 528. | Inpp5j | -2.59 | 0.0062 | Down |
| 529. | Ccl12 | -2.62 | 0.0117 | Down |
| 530. | Slco4c1 | -2.64 | 0.0536 | Down |
| 531. | Klra8 | -2.67 | 0.0557 | Down |
| 532. | Gzma | -2.68 | 0.0000 | Down |
| 533. | Mx1 | -2.68 | 0.0163 | Down |
| 534. | Ifitm6 | -2.70 | 0.0001 | Down |
| 535. | Prss34 | -2.71 | 0.0133 | Down |
| 536. | Gzmd | -2.73 | 0.0059 | Down |
| 537. | Gzmf | -2.74 | 0.0446 | Down |
| 538. | Dmkn | -2.81 | 0.0362 | Down |
| 539. | Tuba8 | -2.83 | 0.0059 | Down |
| 540. | Slc6a4 | -2.86 | 0.0235 | Down |
| 541. | Prg2 | -2.86 | 0.0415 | Down |
| 542. | Retnla | -3.00 | 0.0234 | Down |
| 543. | Vmn2r26 | -3.00 | 0.0261 | Down |
| 544. | Il13 | -3.02 | 0.0427 | Down |
| 545. | Il27 | -3.05 | 0.0161 | Down |
| 546. | Fmo2 | -3.12 | 0.0704 | Down |
| 547. | Alox15 | -3.17 | 0.0108 | Down |
| 548. | Dpep2 | -3.27 | 0.0117 | Down |
| 549. | Gbp2b | -3.35 | 0.0350 | Down |
| 550. | Ear6 | -3.37 | 0.0791 | Down |
| 551. | Acta1 | -3.53 | 0.0356 | Down |
| 552. | Smpd3 | -3.57 | 0.0030 | Down |
| 553. | Cma2 | -3.68 | 0.0803 | Down |
| 554. | Mucl1 | -3.99 | 0.0633 | Down |
| 555. | Gzmg | -5.54 | 0.0001 | Down |

Supplementary table 6: Myeloid / macrophage related pathways at day 30 post tumor challenge

| No. | Description | Set Size | Enrichment Score | NES | p[illegible] |
|---|---|---|---|---|---|
| 1 | GOBP_MYELOID_CELL_ACTIVATION_INVOLVED_IN_IMMUNE_RESPONSE | 95 | -0.773 | -2.679 | 1[illegible] |
| 2 | GOBP_MACROPHAGE_CYTOKINE_PRODUCTION | 38 | -0.758 | -2.261 | 4.[illegible] |
| 3 | GOBP_MACROPHAGE_ACTIVATION | 102 | -0.635 | -2.237 | 1.[illegible] |
